# Tissue-Like Multicellular Development Triggered by Mechanical Compression in Archaea

**DOI:** 10.1101/2024.10.19.619234

**Authors:** Theopi Rados, Olivia S. Leland, Pedro Escudeiro, John Mallon, Katherine Andre, Ido Caspy, Andriko von Kügelgen, Elad Stolovicki, Sinead Nguyen, Inés Lucía Patop, Thiberio Rangel, Sebastian Kadener, Lars D. Renner, Vera Thiel, Yoav Soen, Tanmay A.M. Bharat, Vikram Alva, Alex Bisson

## Abstract

The advent of clonal multicellularity is a critical evolutionary milestone, seen often in eukaryotes, rarely in bacteria, and only once observed in archaea. We show that uniaxial compression induces clonal multicellularity in haloarchaea, forming tissue-like structures. These archaeal tissues are mechanically and molecularly distinct from their unicellular lifestyle, mimicking several eukaryotic features. Archaeal tissues undergo a multinucleate stage followed by tubulin-independent cellularization, orchestrated by active membrane tension at a critical cell size. After cellularization, tissue junction elasticity become akin to animal tissues and give rise to two cell types—radial peribasal and central apicobasal cells— with distinct actin and protein glycosylation polarity patterns. Our findings highlight the potential convergent evolution of a biophysical mechanism in the emergence of multicellular systems across domains of life.

## Main Text

Multicellularity has evolved multiple times across the tree of life, fundamentally reshaping Earth’s biosphere. Comparative studies across these independent transitions have revealed common selective benefits, including increased size, enhanced mechanical strength, and cellular differentiation—principles later confirmed through experimental evolution (1–4). Although well-documented in eukaryotes and prokaryotes, it remains unclear to what extend clonal multicellularity could contribute to the emergence of structural and functional complexity in bacteria and archaea.

Once mistaken for bacteria due to their lack of nuclei, archaea are now recognized as forming a monophyletic group with eukaryotes (5). At the cellular level, most archaea lack a rigid cell wall and are encapsulated by a proteinaceous surface monolayer (S-layer), a 2D paracrystalline lattice composed of glycoproteins (6, 7). While the archaeal envelope structure is thought to make cells mechanically vulnerable, it also facilitates close interactions between cells, such as cell-cell contact and fusion, which may have played a role in the emergence of eukaryotes (8, 9). However, the evolution of mechanosensory responses driven by the lack of a rigid cell wall remains elusive due to the scarcity of in vivo studies. The unique combination of genetic and biophysical traits prompted us to investigate the mechanobiology of archaeal cells, leading to the serendipitous discovery of a reversible, clonal tissue-like multicellular developmental program.

### Uniaxial compression gives rise to clonal, tissue-like multicellularity

To gain insights into the mechanobiology of archaeal cells, we performed confinement experiments with the salt-loving Haloferax volcanii (Hvo), leveraging its straightforward cultivation and genetics (10). First, we established a baseline for mechanically unperturbed haloarchaeal cells trapped within ArcCell, a custom microfluidic device (Fig. 1A and S1A). Cells growing in ArcCell showed cell morphologies comparable to those in bulk liquid cultures, indicating that cells were not mechanically stressed (Fig. 1B and S1B; Movie S1). Next, we imaged cells under agarose pads, a standard technique for microbial immobilization (Fig. 1C). Unlike in ArcCell, agarose pads deformed cells within a single generation (∼2.5 hours) at the lowest agarose concentrations, making pad immobilization incompatible with haloarchaeal prolonged imaging (Fig. S1B, Movie S1). To quantify the compressive forces involved in deforming cells, we measured the pad resistance to mechanical deformation using dynamic mechanical analysis (DMA) (11). The storage moduli of 0.25-3.5% pads revealed that resistance forces of approximately ∼10 kPa are sufficient to deform Hvo cells (Fig. S1C). These values suggest Hvo cells may have viscoelastic properties close to eukaryotic cells such as amoeba and mammalian cells but orders of magnitude lower than most cell-walled organisms (12, 13).

**Fig. 1.**
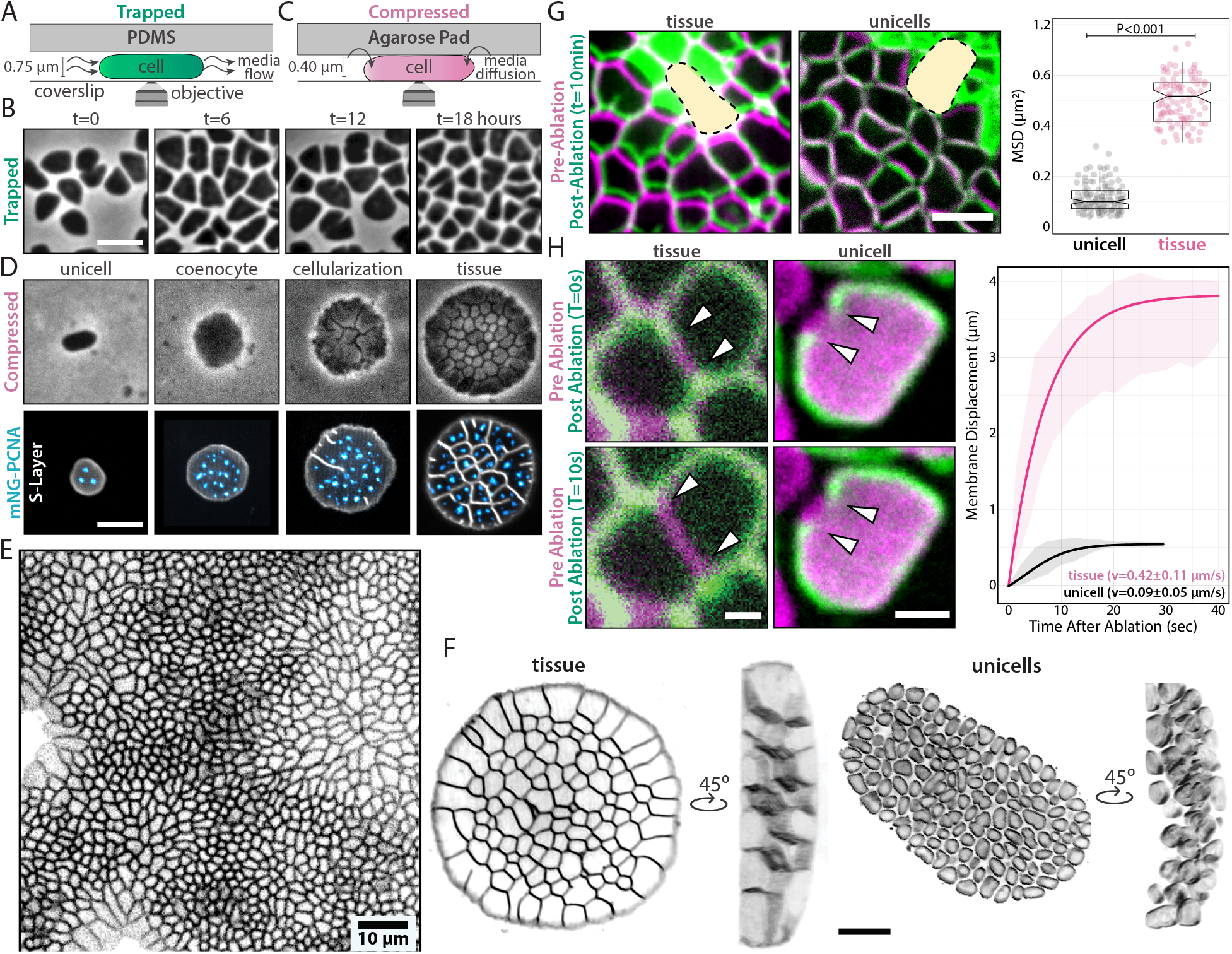
Uniaxial compression triggers multicellular development in *Hfx. volcani*. Schematic of **(A)** microfluidics and **(B)** agarose pad immobilization. Phase-contrast time-lapses of **(C)** trapped and **(D)** compressed cells across ∼6 generations where msfGFP-PCNA foci (blue) represent replication sites. **(E)** Stretched and compressed areas comprised in large monolayers of epithelia-like tissues. **(F)** 3D-SoRa microscopy images of a tissue (top) and unicells (bottom). **(G)** Laser ablation of tissue regions. False-colored overlays of tissues before (magenta) and 10 minutes after (green) ablation. Yellow areas indicate the ablated area. Directional motion from cells was calculated from MSD curves. **(H)** Laser ablation of cell membranes. False-colored overlays of tissues and cells before (magenta) and after (green) ablation. White arrowheads indicate the membrane recoil retraction. Unless specified, scale bars represent 2 μm.

Given the mechanical sensitivity of haloarchaeal cells under pads, we tested the responses of Hvo to compressive forces above 100 kPa, closer to their natural habitat such as the human gut and salt pounds (14–16). Following compression under pads with agarose concentrations 1.5% and higher, cells stopped dividing but continued to grow (Fig. 1D, 2nd panel). After ∼12 hours, cellularization occurred via multiple simultaneous septation events (Fig. 1D, 3rd panel). Finally, cells assembled an epithelial-like monolayers (Fig. 1E; Movie S2), forming connected cellular structures akin to the tessellation patterns in leaf tissues (17).

The observed morphological development before cellularization also resembles the coenocytic phase of the chytrids and chicken embryogenesis, where cells multiply their nuclei without cell division (18, 19). To confirm that these cells are coenocytes, we inquired whether DNA replication progresses during cell division arrest. Time-lapse imaging of cells expressing a fluorescent fusion of the DNA sliding clamp (mNeonGreen-PCNA) showed the multiplication of replication sites during development, confirming that cells under compression go through a coenocytic-like phase preceding cellularization (Fig. 1D; Movie S3).

Interestingly, tissue formation is independent of gravity (Fig. S2A), pad mass or thickness (Fig. S2B), and consistently initiated at the same coenocytic area regardless of pad density (Fig. S2C). These consistent features suggests that archaeal tissue development represents an evolved biological program selected under confinement. Animal and plant cells often sense and activate biochemical pathways in response to surface curvatures and material properties (20, 21). To test the influence of pad’s stiffness, Hvo cells were immobilized under and on top of the same 2.5% agarose pad. After 24 hours, tissue-like structures were exclusively observed under the pads (Fig. S3A), suggesting stiffness alone is not sufficient to induce multicellularity. Furthermore, cells compressed by bilayer “cake” pads developed into tissue-like structures when in contact with the 2.5% agarose surface but remained unicellular under the 0.25% layer (Fig. S3B). Finally, we also observed tissue development only when cells were compressed between two stiff agarose layers, ruled out a specific role of the coverslip other than providing a rigid surface for compression (Fig. S3C). While a direct role for surface stiffness cannot be excluded, we conclude that compression is required for multicellularity.

To determine if cells within multicellular structures retain their S-layer lattice, we cryo-fixed cell resuspension archaeal tissues subjected to mechanical shear and imaged individual cells by electron cryotomography (cryo-ET) (Fig. S4A). Concomitantly, we imaged live cells stained with Brilliant Blue, a fluorescent dye that specifically labels S-layer glycoproteins but not the cytoplasmic membrane (Fig. S4B). Both methods consistently showed S-layer, confirming that archaeal tissues preserve the S-layer in their intercellular spaces. Next, we explored the possibility that tissues arise from cell compaction. 3D imaging revealed extracellular spaces between most unicells but none within tissues (Fig. 1F; Movie S4).

To probe physical connectivity, we used laser ablation to wound areas at the center of the tissues. Cells physically connected by junctions are expected to be collectively pulled towards the center of the injured area. Following ablation, we observed directional movement of cells toward the wounds in tissues, but not in unicells, at speeds of 0.62±0.27 μm/min (Fig. 1G and S4C; Movie S5), similar to those seen in wounded animal tissues, which can vary between 0.2-1.0 μm (22, 23). Given that archaea lack canonical cytoskeleton motors like myosin (24), it is unlikely that this directional motion is a product of active wound healing as mechanistically described in animal tissues. Instead, this synchronic cell migration suggests that both archaeal and animal tissues have similar membrane viscoelastic properties. To test this idea, we measured the retraction rates after ablating the cell envelope in unicells and archaeal tissues. After severing, we observed a membrane recoil in archaeal tissues (0.42±0.11μm/s) (Fig. 1H; Movie S6) similar to those reported in animal tissues (∼0.3μm/s) (25). The apparent higher membrane tension in archaeal tissues compared to compacted unicells (0.09±0.05 μm/s) implies the presence of junctional load bearing structures, placing archaeal tissues as a unique class of multicellularity, exhibiting material properties typical of eukaryotic tissues.

### Archaeal tissues are widespread in Haloarchaea and counter-correlate with biofilm production

To understand the evolutionary origins and diversity of haloarchaeal tissues, we constructed a phylogenomic tree spanning 57 genera, representing all haloarchaeal orders (Data S1). Based on the clade distributions, we cultivated and imaged compressed cells from 52 species across 14 different genera (Fig. S5). Regardless of size or growth rate (Fig. S6A), 61.6% of tested haloarchaeal species formed tissues, with at least one case of tissue development or no multicellularity in every tested genus. Among species that did not form tissue, we identified instances of cell death (Fig. S6B), shape deformation (Fig. S6C), or unnoticeable shape deformations under pads (Fig. S6D). Surprisingly, we observed 3 species – Htg. salina, Ncc. jeotgali, and Hka. jeotgali – that exhibit aggregative multicellularity similar to Methanosarcina, previously the only known multicellular archaeon (Fig. S6E) (26). These results suggest that archaeal tissues emerged early in haloarchaeal evolution, remain dominant in the sampled diversity, and may have been lost multiple times.

Next, we focused on strains from the Haloferax genus, where most species developed tissues (Fig. 2A), with Hfx. prahovense producing larger, deformed tissues (Fig. S7A). In contrast, Hfx. mediterranei (Hmed) and Hfx. gibbonsii (Hgib) were the only strains that failed to develop tissues, growing instead as stacked cell colonies (Fig. 2B and S7A; Movie S7). Although not constituting a monophyletic branch, all three taxa are closely related. Hmed, in particular, is placed on a long branch, suggesting extended time for adaptation following loss or gain of genetic material. We then tested if Hmed cells could still form tissues under higher compression by 5% agarose pads. Hmed cells grew larger than Hvo coenocytes until they split and swarmed outwards (Fig. 2B; Movie S7). Since Hmed’s swarming-like motion resembles bacterial biofilm-dependent gliding (27), we questioned if Hmed cells employ their extracellular matrix to survive compression. Supporting this hypothesis, the relative biofilm mass levels in Hmed were at least twice as high as those in other Hfx strains (Fig. S7B). It remains unclear whether Hmed still hosts the molecular machinery required for multicellularity with mutations suppressing their expression or if it lost the genetic pathways required for multicellularity. The diversification of tissue architecture and survival strategies suggest commonly shared multicellularity with occasional losses.

**Fig. 2.**
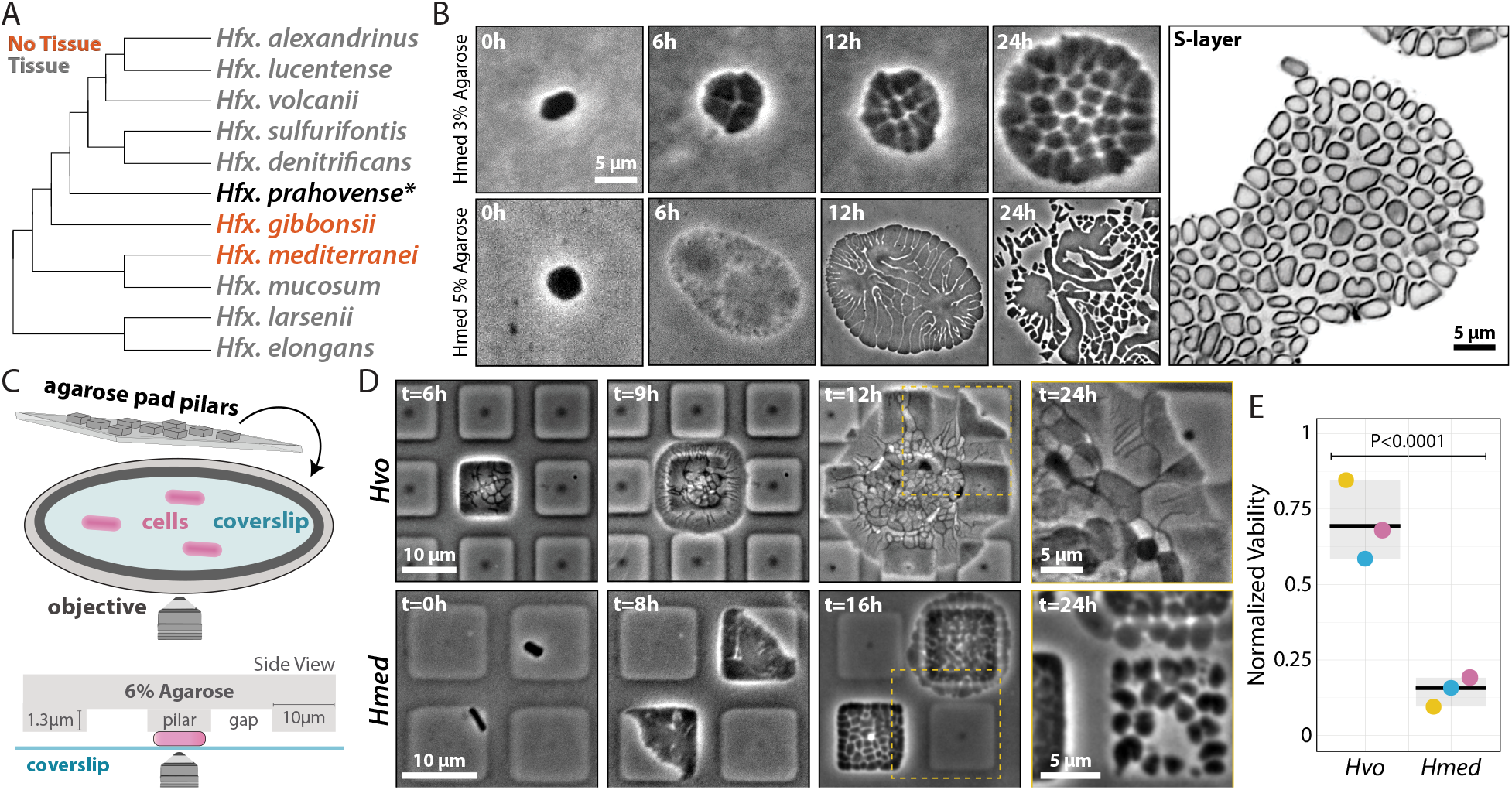
*Hfx. mediterranei* does not form tissues under compression. **(A)** Cladogram depicting evolutionary relationships between compressed Haloferax species. Gray- and red-labeled species represent cells that form or do not form tissues, respectively. Hfx. prahovense is marked with an asterisk as it develops to considerably larger, deformed tissues. For a comprehensive phylogenetic tree, see Supplementary Data S1. **(B)** Phase-contrast time-lapses (left) of Hmed growth under 3% (top) and 5% (bottom) agarose pads. (right) SoRa microscopy of Hmed growth after 24 hours under 3% agarose pads. **(C)** Cartoon representation of microfabricated pillars used in the intermittent compression experiments. **(D)** Phase-contrast time-lapses of Hvo (top) and Hmed (bottom) under micropillar devices. The 24-hour datapoint is represented as a zoom inlet from the yellow area in the previous timepoint. **(E)** Viability of Hvo tissues compared to Hmed cells under micropillars measured by colony formation unit (CFU). Hvo and Hmed viabilities were normalized by their respective liquid cultures.

Despite sharing many similar “weed-like” traits observed in lab, Hmed is still outcompeted in nature by other haloarchaea (28). To test if Hmed’s lack of multicellularity affects its fitness, we compressed cells under microfabricated pillars intercalated with “relief” zones (Fig. 2C). This setup allowed us to observe how these cells navigated mechanical “escape room” challenges, mimicking their natural habitat. While Hvo cells managed to propagate even with low initial cell numbers, Hmed showed a 4.3-fold decrease in viability compared to Hvo under pillars (Fig. 2D and E; Movie S8). In contrast, Hvo tissues showed only a 1.8-fold decrease in viability compared to unicells from liquid cultures, and 1.4-fold decrease compared to Hvo colonies from agar plate streaks (Fig. S7C).

The survival rates between Hvo and Hmed suggests that tissues can revert to unicells. To observe this transition under the microscope, we shear-shocked tissues by injecting liquid media in parallel to pads. As a result, cells detached from tissues and transitioned to motile rods, swimming away from the initial compression zones (Movie S9). Our findings suggest that mechano-responsive multicellularity is an adaptive trait and likely beneficial in compressive zones exceeding lethal thresholds in haloarchaeal environments, such as desiccated salt plates, animal guts, and microbial biofilms (29).

### Archaeal tissues show radial symmetry with distinct cell types and undergo FtsZ-independent cellularization

Time-lapses of Hvo experiencing intermittent compression revealed tissues with larger peripheral cells compared to those under pads (Fig. 2D). Changes in cell size and shape could result either from the uneven distribution of mechanical forces within the device or from mechanosensation by specific cells. Since specialized cell types are a hallmark of complex multicellularity (30), we characterized the cellular morphology and growth in different tissue regions. 3D-STED micrographs showed two cell profiles: wider and shorter at the periphery, and taller at the center of tissues (Fig. 3A, Movie S10). Moreover, peripheral cells are not in contact with the surface of pads, indicating their low height is not a product of compression. In contrast, central cells are in physical contact with compression areas, implicating they are directly responding to the mechanical compression from pads. To confirm that peripheral cells do not interact with compression zones, we repurposed a method inspired by traction force microscopy (31). Imaging fluorescent beads adhered to pads, we observed a vertical upward displacement of beads over the tissues’ central regions, but not over their periphery (Fig. S8A). Based on their radial symmetry and position within the tissue, we named tissue cells as peribasal (peripherical basal regions) and apicobasal (central apical and basal regions) (Fig. 3B).

**Fig. 3.**
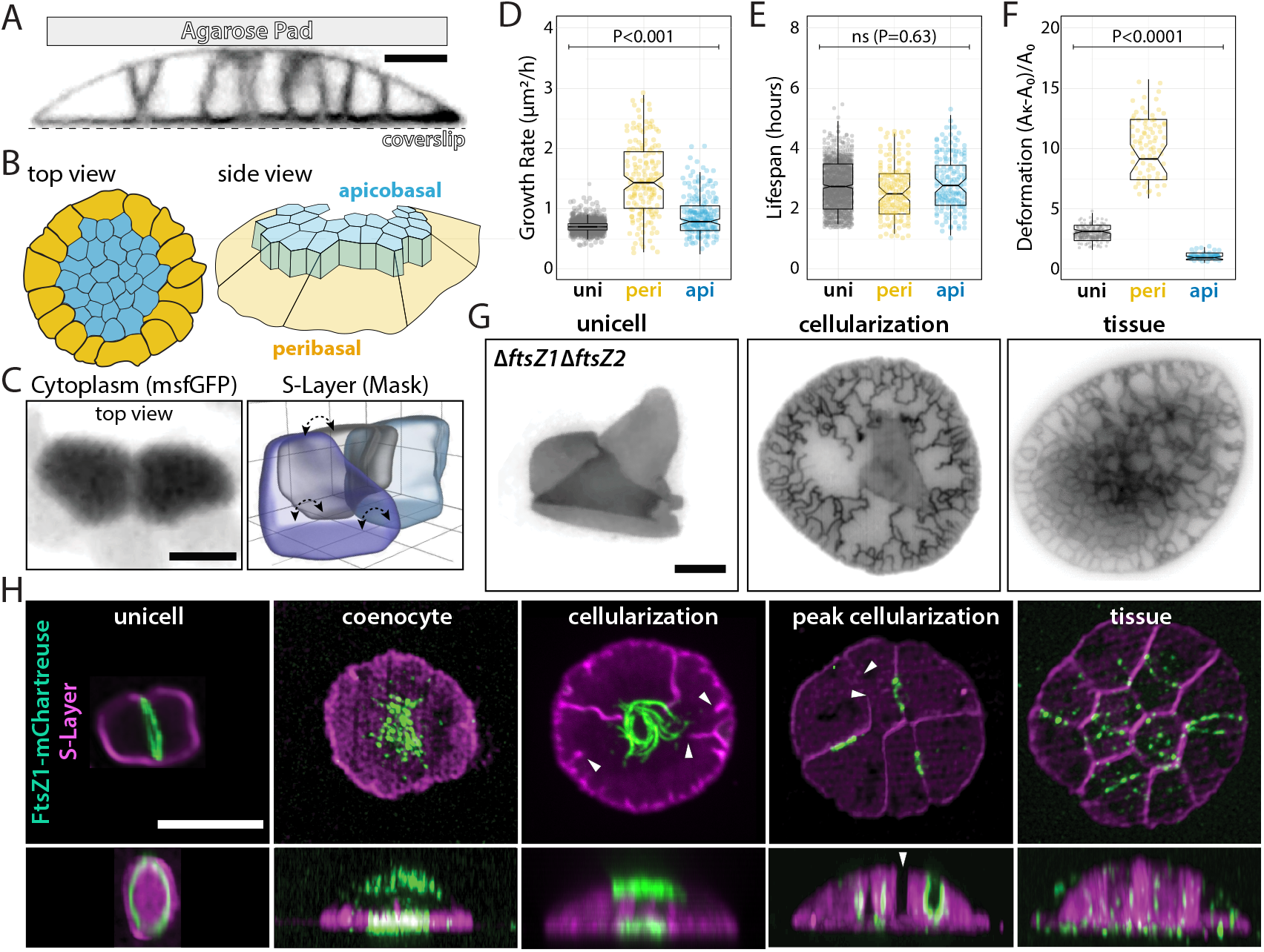
Cellularization is independent of FtsZs and results in two cell types. **(A)** 3D-STED super-resolution microscopy of the cellularization process. **(B)** Cartoon representation of top and side views showing apicobasal and peribasal cell types within tissues. **(C)** (Left) iSIM of cells expressing cytoplasmic msfGFP and (right) 3D outline masks of scutoid apicobasal cells segmented from 3D-STED highlights apicobasal scutoid cells. Dashed arrows indicate the different apicobasal surface neighbors across apicobasal planes. **(D-F)** Cell area growth rate **(D)**, lifespan **(E)** and deformation **(F)** measurements of unicells, peribasal, and apicobasal cells under 2.5% agarose pads from phase-contrast time-lapses. **(G)** Epifluorescence microscopy of representative cell-division impaired ΔftsZ1ΔftsZ2 cells across different developmental stages. Early cellularization represents cells that just entered the cellularization stage. Peak cellularization represents cells at the onset of completing cellularization. **(H)** 3D-SoRa microscopy of representative cells expressing FtsZ1-mChartreuse across different developmental stages. White arrowheads indicate septation sites without FtsZ1 signal. Scale bars: 2 μm.

From our 3D-STED projections, we observed that the apicobasal cell shape resembles irregular 3D-polygonal scutoids that stabilize curved epithelia during embryogenesis (32). Scutoid cells maximize tissue packing by intercalating along their height, resulting in varying neighbor counts across apicobasal planes. From 3D outline segmented masks, we observed variation in cell-cell neighborhoods across the apicobasal regions (Fig. 3C). Another important feature of animal scutoid cells is minimizing energy by distributing membrane tension. This led us to determine if archaeal scutoids maintain a constant surface-to-volume ratio compared to peribasal cells. Although peribasal volumes and surface areas were larger than apicobasal cells, their ratios remained consistent (Fig. S8B). Our data suggests that scutoid-like cells may have emerged early, predating eukaryotes.

Mechanically stressed cells are typically smaller, grow slower, and have shorter lifespans compared to unstressed cells (33). Thus, to determine whether apicobasal and peribasal shapes result from physical packing, we compared their growth rates and lifespans to unicells. Surprisingly, both cell types exhibited higher growth rates (Fig. 3D) and similar lifespans (Fig. 3E) relative to unicells. Altogether, our data supports the conclusion that tissue morphology cannot be summarized by simple mechanical or nutritional stresses.

To expand on the apicobasal and peribasal specialization, we examined if differences in size and shape could translate to distinct viscoelastic properties. After breaking tissues by mechanical shear, we subjected individual cells to another compression cycle and measured their deformation. While peribasals deformed ∼3-fold more than unicells, apicobasals were ∼2.5 more rigid than (Fig. 3F, Movie S11). This supports a model where stiffer apicobasal cells counteract pad compression so flexible peribasal cells can crawl out of compression zones.

Specialized cell types in animal tissues often adapt their cytoskeleton across different mechanical cues. To test if Hvo tissues still undergo cytokinesis by unicellular mechanisms, we examined the role of tubulin paralogs FtsZ1 and FtsZ2. Although FtsZs are required for cell division in unicells (34), the ΔftsZ1ΔftsZ2 mutant was still capable of cellularization under compression with “zig-zag” askew cell junctions (Fig. 3G). These observations parallel some animals and multicellular holozoans, where microtubules are dispensable (35). To further understand the role of FtsZs during development, we created a functional knock-in FtsZ1-mChartreuse fusion (36) (Fig. S9A). In unicells, FtsZ1 formed continuous Z-rings, converting to scattered FtsZ1 filaments at the top and bottom of the central coenocytic region (Fig. 3H). Supporting the conclusion that cellularization is FtsZ-independent, FtsZ1 was absent at 42% of the junction furrows (Fig. S9B, white arrowheads). Collectively, these observations support a sequential developmental program, with tissues relying on distinct molecular machinery for cell division that is absent in unicells.

### Archaeal tissue cellularization is triggered by envelope tension

Evidence from flies and unicellular holozoans suggests that the timing of cellularization correlates with the ratio between the number of nuclei and cell volume (N/C) (37). To test if N/C is involved in triggering cellularization in coenocytes, we imaged cells expressing mNeonGreen-PCNA. We quantified the number of replication sites and the fluorescence intensity within replication sites relative to cell area as a proxy for N/C. If N/C were critical for cellularization, DNA replication would need to increase at a faster rate than cell area. However, replication rates remained constant relative to cell area indicating that N/C does not influence cellularization (Fig. S10A).

Next, we tested if other markers, such as size, time, or the amount of added cell volume, coincide with cellularization (38). By tracking these single-cell parameters at cellularization (Fig. 4A and 4B), we observed a lower coefficient of variance for coenocyte area (CV=9.6%) compared to area added and time (CV=24.2% and 47.1%, respectively) (Fig. 4C), suggesting that cellularization happens at a specific coenocyte size. To validate this correlation, we tracked the development of compressed ΔftsZ2 cells. Because ΔftsZ2 cells do not divide when growing exponentially in liquid cultures, they are larger than control cells and would be expected to cellularize sooner. Indeed, ΔftsZ2 cells cellularized approximately 4.2 times faster and added 3.7 times less area but at a similar area threshold to wild type (Fig. 4C).

**Fig. 4.**
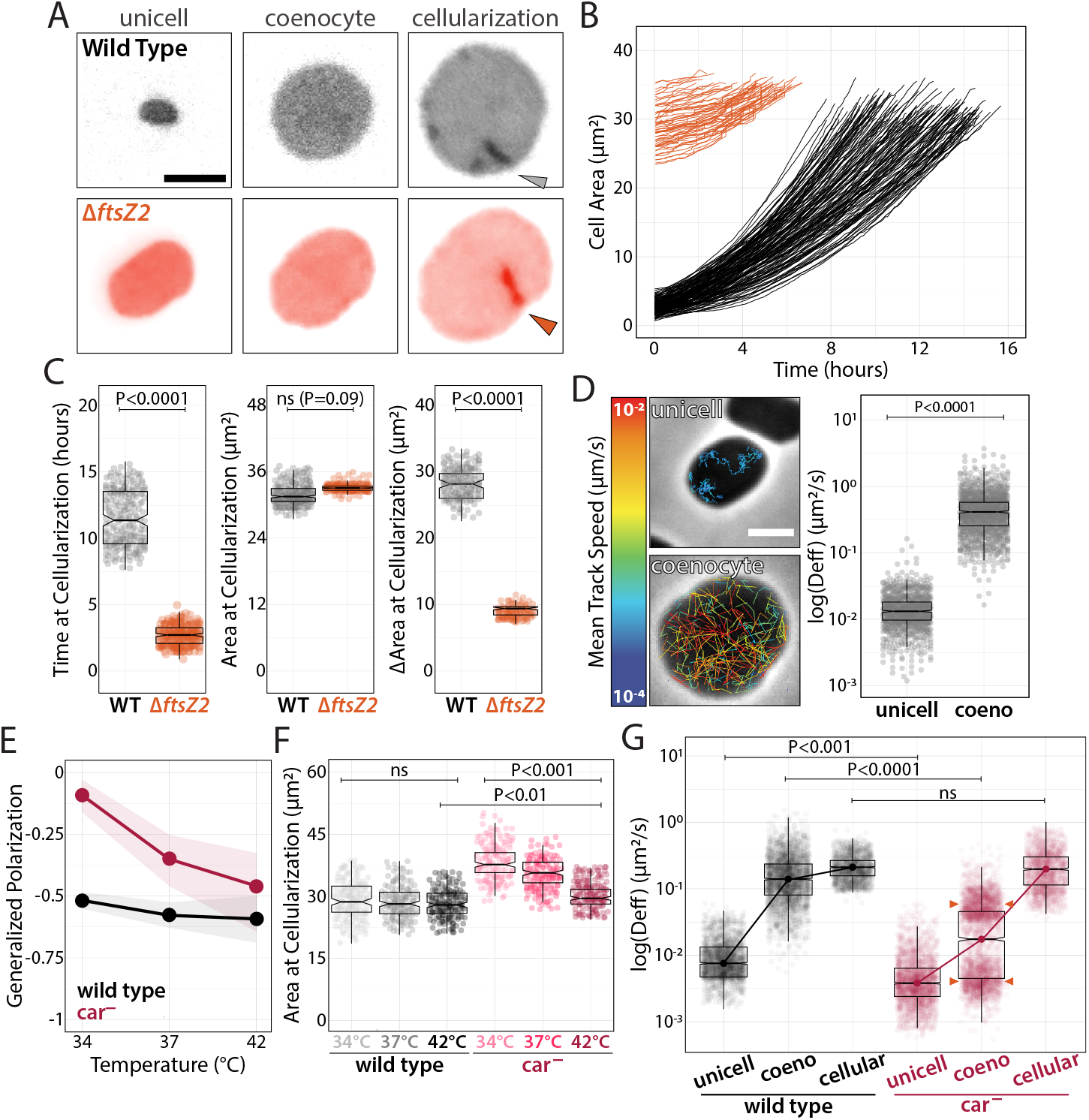
Tissue cellularization is triggered by coenocytic size through a membrane tension threshold. **(A)** Epifluorescence micrographs of wild-type (top) and ΔftsZ2 (bottom) cells from compression to cellularization. (B) Single-cell growth curves from compression to cellulation onset. (C) Time, cell area, and cell area added at the onset of cellularization. (D) Phase-contrast images of unicell and coenocytes and bSpoJ single-molecule tracks overlays false-colored relative to their mean speed. Effective diffusion coefficients calculated from MSD curves. (E) Live-cell Generalized Polarization measurements of wild-type and car-stained with Laurdan. (F) Area at cellularization measurements of wild-type and car-cells from phase-contrast time-lapses across temperatures. (G) Wild type and car-membrane fluidity calculated by bSPoJ effective diffusion coefficients across developmental stages at 34°C. Correlation between cellularization areas and bSpoJ diffusion in wild-type and car-cells at 34°C. Scale bars: 2 μm.

Although cell size is consistent at cellularization, the underlying mechanism remains unclear. Given the differences in membrane tension between tissues and unicells, we hypothesized that cell envelopes accumulate elastic strain as coenocytes become larger, reaching a critical threshold to initiate cellularization. To measure the real-time membrane tension in coenocytes, we created bSpoJ, a single-molecule live-cell membrane fluidity reporter. bSpoJ is a chimeric transmembrane domain of SpoIIIJ from Bacillus subtilis, with a secretion signal peptide from Hvo, and HaloTag. Particle tracking of bSpoJ in Hvo showed a 30.3-fold increase in apparent diffusion in coenocytes compared to unicells, an order of magnitude above the 2.5-fold increase in cell area (Fig. 4D and S10B; Movie S12 and S13).

The disproportionate increase in membrane tension relative to cell area suggests that the biochemical composition of the coenocytes actively changes throughout development. To test this hypothesis, we inquired if perturbing membrane fluidity as a proxy for tension would affect the cell area at cellularization. Because carotenoids are thought to play a similar role in bacteria as cholesterol in eukaryotes (39), we tested if membrane fluidity would fluctuate in cells unable to produce carotenoids (car−). As an orthogonal approach to bSpoJ, we measured membrane fluidity in cells stained with Laurdan, a fluorescent probe, which intercalates between phospholipids and reports generalized polarization (GP) (40). Despite the lack of obvious phenotypic defects in car− unicells at higher or lower temperatures (Fig. S10C), car− showed a weaker membrane fluidity homeostasis compared to wild type when cultivated at lower temperatures (Fig. 4E and S10D). Consistent with this, car− cells showed in a decrease of coenocyte viability (Fig. S10E). The colony pigmentation also appeared to be attenuated in Hmed and other Hfx species that show defective tissue structures (Fig. S10F and S11), suggesting a positive correlation between carotenoids and tissue development, in opposition to biofilm production.

The loss of car− membrane fluidity control implies that coenocytes fail to trigger cellularization below critical tension thresholds. Hence, we expect changes in its cell size at, but survival cells must reach the same membrane tension threshold as in the wild type. As predicted, we observed an increase in cell area at cellularization correlated to car− loss of viability at lower temperatures (Fig. 4F), suggesting car− cells can only match membrane tension threshold at larger cell areas. To directly probe membrane fluidity at cellularization, we compared bSpoJ diffusion between wild-type and car− cells at 34°C (Fig. 4G). Once again, we observed lower membrane fluidity in car− unicells compared to wild type as recorded by generalized polarization. Above all, bSpoJ diffusion at tissue cellularization were similar in wild type and car−, supporting a tension control mechanism. Curiously, we observed a bimodal distribution in car− coenocytes (orange arrowheads). We speculate that one sub-population with higher bSpoJ diffusion rates reach the membrane tension cutoff, but still lower membrane tension compared to wild type, therefore cellularize at a larger size. In contrast, the second group stochastically fails to satisfy the tension-area ratio requirement and undergo lysis. Altogether, these results highlight a mechanoresponsive mechanism in haloarchaea, wherein cells actively regulate membrane fluidity and developmental progression to prevent premature or delayed cellularization.

### Actin and N-glycosylation are fiducial markers for archaeal tissue polarity

Next, we searched for developmental factors specific to multicellular lifestyle by performing RNA-seq on each developmental stage (Fig. 5A). Gene expression profiles revealed that coenocytes repress twice as many as upregulated genes, possibly involving pathways related to stress response and cell division arrest (Fig. S9A). Curiously, although the number of differentially expressed genes (∼28% of the genome) are distributed equally across the genome (Fig. S9B), we observed significant enrichment in the expression of genes located in the third quarter of the genome in tissues, suggesting the existence of possible multicellular gene islands (Fig. S9C). From hundreds of candidate genes, we focused on four main classes: cell surface biogenesis, cytoskeleton and mechanosensing, signaling, and cell cycle (Data S2).

**Fig. 5.**
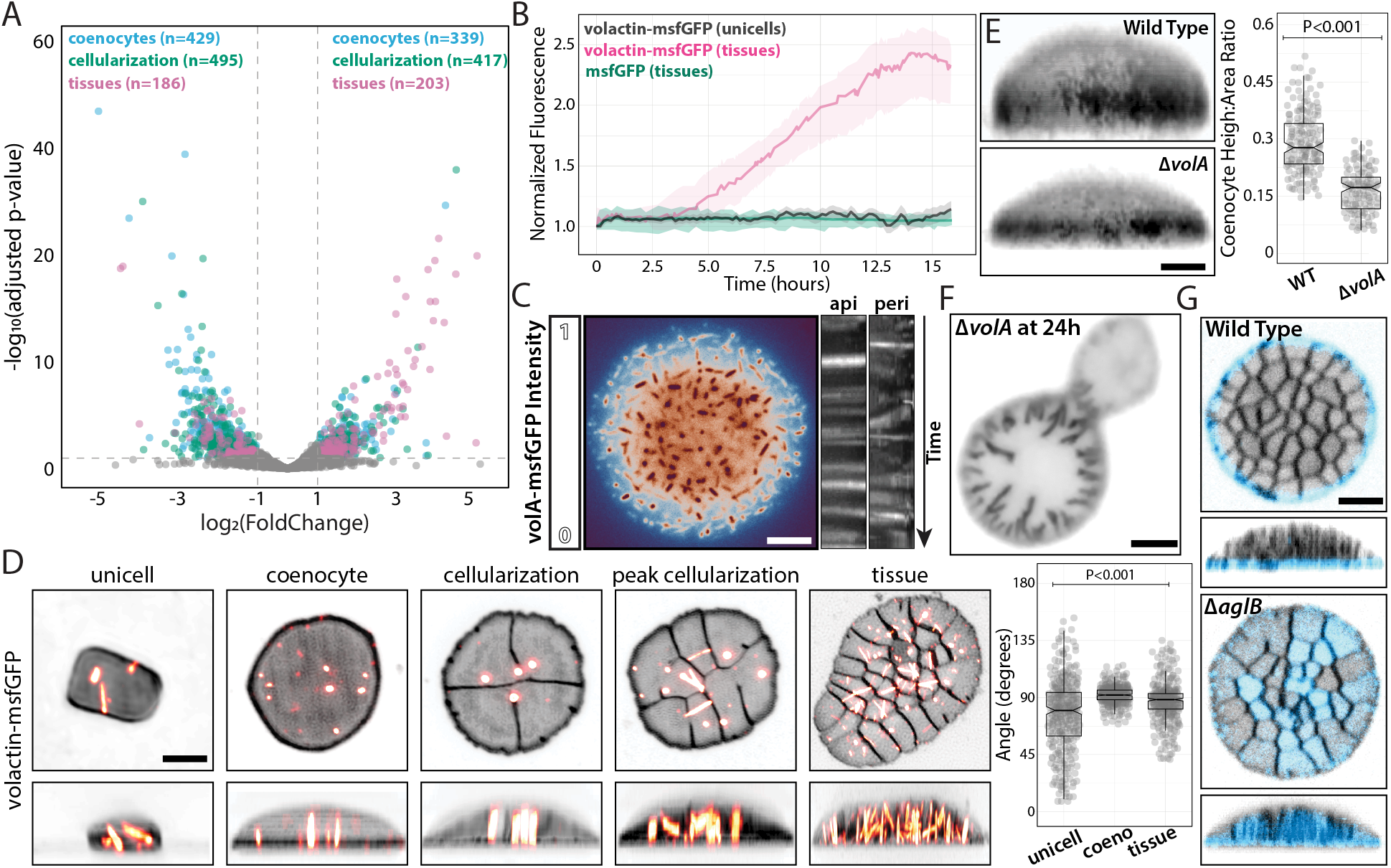
Volactin and N-glycosylation are tissue-specific polarity markers. **(A)** Volcano plot overlays from RNA-seq datasets collected across developmental stages and normalized by liquid unicellular cultures. Parentheses indicate the number of candidates above the arbitrary cutoff. **(B)** Normalized fluorescence by cell area of volA-msfGFP and constitutively expressed cytoplasmic msfGFP from confocal time-lapses. **(C)** Epifluorescence micrograph of a false-colored tissue relative to volA-msfGFP fluorescence (left) and dynamics of volA-msfGFP polymers represented by kymographs (right) from peribasal and apicobasal regions. **(D)** 3D-SoRa projections of representative cells expressing volA-msfGFP across developmental stages (left) and volA cable angle measurements relative to the coverslip plane (right). **(E)** height measurements of wild-type and ΔvolA coenocytes from 3D-SoRa projections. **(F)** Representative epifluorescence micrograph of a ΔvolA cell stalled at cellularization. **(G)** 3D-confocal projections of cell surface N-glycosylated proteins in wild-type (top) and ΔaglB (bottom) tissues stained by ConA-Alexa488. Scale bars: 2 μm.

Given the importance of actin in eukaryotic tissue biogenesis, we investigated the role of volactin (volA), the only identified actin homolog in *Hvo* (*39*), which was upregulated during cellularization and tissues in our RNA-seq datasets. Time-lapses of developing cells showed increased volA-msfGFP signal, consistent with our RNA-seq dataset, reaching a steady-state at the end of cellularization (Fig. 5B; Movie S12). In contrast, trapped unicells showed unaltered volA-msfGFP expression, as did tissues expressing cytoplasmic msfGFP. Moreover, volactin polymers were less abundant but highly dynamic in uncompressed apical cells compared to unicells and basal scutoids (Fig. 5C; Movie S13). Notably, cells displayed different volA spatiotemporal patterns during development, with volactin cables aligned in parallel in coenocytes and apical cells within tissues (Fig. 5D; Movie S14). Compression of Δ*volA* cells resulted in shorter coenocytes than wild type (Fig. 5E) and delayed or stalled tissue maturation (Fig. 5F). VolA and other factors have recently been implicated in regulating the development of rod-shaped (motile) and disk-shaped (sessile) cell types (*39, 40*). Therefore, based on cellular localization, dynamics, and morphological phenotypes, we propose that volactin may mirror eukaryotic actin’ s multifunctional roles as a mechanosensing cytoskeleton and polarity factor in tissues (*41*).

Leveraging the differences in tissue formation across *Hfx* species (Fig. 2A), we used comparative genomics and over-representation analysis to create datasets contrasting the presence and absence of orthologous protein groups (orthogroups) among (i) *Hvo* and *Hmed* (Fig. S9D; Data S3); (ii) 4 *Hfx* species (Fig. S9E; Data S4); and (iii) orthogroups enriched in species that form tissues (Data S5). We found a strong representation of orthogroups related to protein glycosylation, sugar metabolism, and transport in all three datasets. Protein glycosylation has historically been a well-studied area in haloarchaea due to its role in mating and envelope biogenesis (*42*). N-glycosylation is also crucial for cell identity, polarity, junctions, and adhesion in multicellular eukaryotes (*43*– *45*). To explore the role of N-glycosylation, we labeled cells with ConA-Alexa488, a fluorescent lectin conjugate impermeable to cells, which has an affinity for various mannose glycol groups. While ConA-Alexa488 did not bind to unicellular colonies or cells in liquid cultures (Fig. S10), it exhibited a radial, halo-like localization at the periphery of apical cells in tissues (Fig. 5G; Movie S15). To test whether these radial patterns are formed by secreted proteins to the extracellular matrix space, we examined Δ*pibD* mutant that cannot process and secrete pili-dependent extracellular proteins, preventing cell adhesion to surfaces (*46*). We observed no differences on ConA-Alexa488 localization or tissue structure in Δ*pibD* tissues, suggesting that surface-adhesion-dependent biofilm is not critical for cell junctions and that ConA-Alexa488 patterns represent cell-type specific glycosylation connected to the outer surface of apical cells. To further test this idea, we imaged tissues of different N-glycosylation mutants, with Δ*aglB* being the only strain to show a disruption in ConA-Alexa488 localization, resulting in binding to both apical and basal surfaces of tissues (Fig. 5G and Fig. S10). Therefore, we propose that AglB, an oligosaccharyltransferase required for S-layer N-glycosylation (*42*), mediate tissue cell-surface polarity. This discovery adds another function to S-layer glycoproteins beyond morphogenesis and mating, establishing their role in tissue polarity.

Our discovery of clonal, tissue-like multicellularity in archaea highlights the potential of archaeal mechanobiology to shed light on the emergence of complexity in nature. Notably, this is not the first developmental program identified in haloarchaea, joining the rod-shaped (motile) and disk-shaped (sessile) shape-shift transitions (*40*). These cell types are connected by volactin, which is also required for disk formation (*39*). The coordinated alignment of volactin cables indicates that actin cables might sense membrane tension, curvature, or mechanically supporting coenocytes and basal cells (*47*). While these scenarios are not mutually exclusive, volA functions in different developmental programs can address a longstanding gap between monofunctional prokaryotic and multifunctional eukaryotic cytoskeletal polymers (*41, 48*). Moreover, the developmental defects observed in Δ*ftsZ1*-*2* tissues underscore the transition from tubulin-dependent cellularization in unicellular prokaryotes to actin-dependent in eukaryotes.

Integrating biophysics and evo-devo (phys-evo-devo) offers a new framework for understanding microbial biofilms and membrane homeostasis, which regulate the transition between different (often contradictory) types of multicellularity. Evolutionarily, our data suggest the need to reinterpret past evidence for the origin of eukaryotic multicellularity, such as fossils identified as holozoans (*49*). Based on their size, shape, and lack of evident eukaryotic features, these fossils may represent ancestors of archaeal tissues. Future studies should aim to uncover the biochemical nature of cell-cell junctions and explore the presence of archaeal tissues in other phyla, particularly those closer to eukaryotes, like Asgard archaea.

## Acknowledgments

The Bisson Lab appreciates Bruce Goode (Brandeis U) for access to the SoRa microscope, and also Avital Rodal (Brandeis U), Buzz Baum (MRC-LMB), and Harold Erickson (Duke U) for constructive suggestions on the manuscript. STED and spectral confocal microscopy were performed at the Brandeis Light Microscopy Core Facility. We want to thank Steven Bruckbauer for the chromosomal mNeonGreen-PCNA strain. TR would like to thank the MBL Woods Hole Physiology Course and Biovis for iSIM use. We are grateful to the Scientific Community Image Forum and the Center for Open Bioimage Analysis (COBA) for fostering an invaluable intellectual community dedicated to training microscopists in image analysis. We acknowledge the MRC LMB electron microscopy facility for sample preparation and data collection. This work was supported by the NIH grant 1S10OD034223-01 for the Abberior Facility Line STED microscope, housed in the Brandeis Light Microscopy Core Facility; Moore–Simons Project on the Origin of the Eukaryotic Cell (doi:10.46714/735929LPI) awarded to AB; the Human Frontiers Science Program (RGY0074/2021) awarded to AB, TAMB, and VA; and the Brandeis National Science Foundation (NSF) Materials Research Science and Engineering Center (MRSEC) Bioinspired Soft Materials (NSF-DMR 2011846). AB is a Pew Scholar in the Biomedical Sciences, supported by The Pew Charitable Trusts. This work was supported by the Medical Research Council, as part of United Kingdom Research and Innovation (also known as UK Research and Innovation) [Programme MC_UP_1201/31 to TAMB]. TAMB would like to thank the European Molecular Biology Organization, the Wellcome Trust (Grant 225317/Z/22/Z) and the Leverhulme Trust. IC was supported by an EMBO Long-Term Fellowship (ALTF 92-2022). This work was partly supported by institutional funds of the Max Planck Society. VA would like to thank Andrei Lupas (MPI Tübingen) for their continued support.

## Author contributions

Conceptualization: TR, OL, AB

Methodology: TR, OL, PE, JM, KA, ES, SN, IP, LTR, LR, YS, AB

Investigation: TR, OL, PE, JM, KA, IC, AvK, SN, LTR, TAMB, AB

Visualization: TR, OL, PE, KA, AB

Funding acquisition: AB, VA, TAMB, VT

Project administration: AB

Supervision: TR, SK, YS, TAMB, VA, AB

Writing – original draft: TR, OL, AB

Writing – review & editing: TR, OL, PE, JM, KA, IC, AvK, ES, SN, IP, SK, LR, VT, YS, TAMB, VA, AB

Conceptualization: SBB, DLA, MPW

## Competing interests

Authors declare that they have no competing interests.

## Data and materials availability

All custom software used in this work is publicly available without restrictions at https://github.com/Archaea-Lab/Multicell_paper and other repositories described in the Supporting Materials. The complete raw RNA-seq datasets presented in this study can be found in NCBI GEO online repository PRJNA1165275.

## Supplementary Materials

Materials and Methods Figs. S1 to S10

Tables S1 to S6

References (50–140)

Movies S1 to S15

## Notes

### Competing Interest Statement

The authors have declared no competing interest.

### Summary of Updates

Inclusion of new data without change in the conclusions: Fig. S2 and S3: Inclusion of pad compression controls, such as titration of pad weight, thickness, and agarose concentration; tissue development against gravity; consistent cell size at cellularization across agarose concentrations above pad stiffness threshold; agarose-agarose pad sandwiches. Fig 1H: Conversion of the cell junction ablation graph from normalized to absolute recoil distances. Fig. 2A: Inclusion of Hfx. gibbonsii as a non-tissue forming organism. Fig. S6A: Inclusion of correlation between unicell cell area and growth across haloarchaeal species, ruling out their role in predicting tissue formation. Fig. 2D: Inclusion of Hmed low-inoculum images and movie (Movie S8) in micropillars. Movie S9: Disassembly of tissue structures and conversion to motile rods after decompression. Fig. 2E and S7C: Viability measurements of Hmed versus Hvo within micropillar devices, and Hvo tissues versus Hvo colonies. Fig. S8A: Inclusion of Traction Force Microscopy experiment to support the claim that peripheral cells are not in contact with pads. Fig. 3B: Change in cell-type terminology. We opted for peribasal (radial cells occupying the periphery and basal regions exclusively) and apicobasal (scutoids occupying both basal and apex regions). Fig. 3C and Movie S10: inclusion of 3D-segmented scutoid cell masks Fig. 3F and Movie S11: Deformation measurements following release and recompression of tissue cells, corroborating cell-typing. Fig. 4G: Membrane fluidity measurements by bSpoJ tracking in Wild Type and car- unicells, coenocytes, and at cellularization supporting the tension-control model.

## References

1. S. Datta, W. C. Ratcliff, Illuminating a new path to multicellularity. Elife 11, e83296 (2022).

2. M. D. Herron, J. M. Borin, J. C. Boswell, J. Walker, I.-C. K. Chen, C. A. Knox, M. Boyd, F. Rosenzweig, W. C. Ratcliff, De novo origins of multicellularity in response to predation. Sci. Rep. 9, 2328 (2019).

3. S. Jacobeen, J. T. Pentz, E. C. Graba, C. G. Brandys, W. C. Ratcliff, P. J. Yunker, Cellular packing, mechanical stress and the evolution of multicellularity. Nat. Phys. 14, 286–290 (2018).

4. T. Brunet, M. Albert, W. Roman, M. C. Coyle, D. C. Spitzer, N. King, A flagellate-to-amoeboid switch in the closest living relatives of animals. Elife 10 (2021).

5. O. Dudin, S. Wielgoss, A. M. New, I. Ruiz-Trillo, Regulation of sedimentation rate shapes the evolution of multicellularity in a close unicellular relative of animals. PLoS Biol. 20, e3001551 (2022).

6. B. Baum, A. Spang, On the origin of the nucleus: a hypothesis. Microbiol. Mol. Biol. Rev. 87, e0018621 (2023).

7. T. A. M. Bharat, A. von Kügelgen, V. Alva, Molecular logic of prokaryotic surface layer structures. Trends Microbiol. 29, 405–415 (2021).

8. M. F. Abdul-Halim, S. Schulze, A. DiLucido, F. Pfeiffer, A. W. Bisson Filho, M. Pohlschroder, Lipid Anchoring of Archaeosortase Substrates and Midcell Growth in Haloarchaea. MBio 11, 10.1128/mbio.00349-20 (2020).

9. D. Moi, S. Nishio, X. Li, C. Valansi, M. Langleib, N. G. Brukman, K. Flyak, C. Dessimoz, D. de Sanctis, K. Tunyasuvunakool, J. Jumper, M. Graña, H. Romero, P. S. Aguilar, L. Jovine, B. Podbilewicz, Discovery of archaeal fusexins homologous to eukaryotic HAP2/GCS1 gamete fusion proteins. Nat. Commun. 13, 3880 (2022).

10. B. Baum, D. A. Baum, The merger that made us. BMC Biol. 18, 72 (2020).

11. M. Pohlschroder, S. Schulze, Haloferax volcanii. Trends Microbiol. 27, 86–87 (2019).

12. K. P. Menard, N. Menard, Dynamic Mechanical Analysis (CRC Press, London, England, ed. 3, 2020.

13. J.-P. Rieu, H. Delanoë-Ayari, S. Takagi, Y. Tanaka, T. Nakagaki, Periodic traction in migrating large amoeba of Physarum polycephalum. J. R. Soc. Interface 12, 20150099 (2015).

14. J. Lee, K. Jha, C. E. Harper, W. Zhang, M. Ramsukh, N. Bouklas, T. Dörr, P. Chen, C. J. Hernandez, Determining the Young’s modulus of the bacterial cell envelope. ACS Biomater. Sci. Eng. 10, 2956–2966 (2024).

15. C. M. Chibani, A. Mahnert, G. Borrel, A. Almeida, A. Werner, J.-F. Brugère, S. Gribaldo, R. D. Finn, R. A. Schmitz, C. Moissl-Eichinger, A catalogue of 1,167 genomes from the human gut archaeome. Nat. Microbiol. 7, 48–61 (2022).

16. C. F. Guimarães, L. Gasperini, A. P. Marques, R. L. Reis, The stiffness of living tissues and its implications for tissue engineering. Nat. Rev. Mater. 5, 351–370 (2020).

17. S. Shad, N. Razaghi, D. Zivar, S. Mellat, Mechanical behavior of salt rocks: A geomechanical model. Petroleum 9, 508–525 (2022).

18. E. Kalinka, S. M. Brody, A. J. M. Swafford, E. M. Medina, L.K. Fritz-Laylin, Genetic transformation of the frog-killing chytrid fungus Batrachochytrium dendrobatidis. Proc. Natl. Acad. Sci. U. S. A. 121, e2317928121 (2024).

19. Y. S. Hwang, M. Seo, H. J. Choi, S. K. Kim, H. Kim, J. Y. Han, The first whole transcriptomic exploration of pre-oviposited early chicken embryos using single and bulked embryonic RNA-sequencing. Gigascience 7, 1–9 (2018).

20. O. Hamant, E. S. Haswell, Life behind the wall: sensing mechanical cues in plants. BMC Biol. 15, 59 (2017).

21. C. Villeneuve, A. Hashmi, I. Ylivinkka, E. Lawson-Keister, Y. A. Miroshnikova, C. Pérez-González, S.-M. Myllymäki, F. Bertillot, B. Yadav, T. Zhang, D. Matic Vignjevic, M. L. Mikkola, M. L. Manning, S. A. Wickström, Mechanical forces across compartments coordinate cell shape and fate transitions to generate tissue architecture. Nat. Cell Biol. 26, 207–218 (2024).

22. D. Kuipers, A. Mehonic, M. Kajita, L. Peter, Y. Fujita, T. Duke, G. Charras, J. E. Gale, Epithelial repair is a two-stage process driven first by dying cells and then by their neighbours. J. Cell Sci. 127, 1229–1241 (2014).

23. W. Kong, O. Loison, P. Chavadimane Shivakumar, E. H. Chan, M. Saadaoui, C. Collinet, P.-F. Lenne, R. Clément, Experimental validation of force inference in epithelia from cell to tissue scale. Sci. Rep. 9, 14647 (2019).

24. L. E. Mayerhofer, A. J. Macario, E. Conway de Macario, Lamina, a novel multicellular form of Methanosarcina mazei S-6. J. Bacteriol. 174, 309–314 (1992).

25. N. Wadhwa, H. C. Berg, Bacterial motility: machinery and mechanisms. Nat. Rev. Microbiol. 20, 161–173 (2022).

26. A. Oren, Why isn’t Haloferax mediterranei more “weed-like”? FEMS Microbiol. Lett. 364 (2017).

27. S. Kim, M. Pochitaloff, G. A. Stooke-Vaughan, O. Campàs, Embryonic tissues as active foams. Nat. Phys. 17, 859–866 (2021).

28. E. P. Bingham, W. C. Ratcliff, A nonadaptive explanation for macroevolutionary patterns in the evolution of complex multicellularity. Proc. Natl. Acad. Sci. U. S. A. 121, e2319840121 (2024).

29. B. E. Shapiro, H. Jonsson, P. Sahlin, M. Heisler, A. Roeder, M. Burl, E. M. Meyerowitz, E. D. Mjolsness, Tessellations and pattern formation in plant growth and development, arXiv [q-bio.CB] (2012). http://arxiv.org/abs/1209.2937.

30. P. Gómez-Gálvez, P. Vicente-Munuera, A. Tagua, C. Forja, A. M. Castro, M. Letrán, A. Valencia-Expósito, C. Grima, M. Bermúdez-Gallardo, Ó. Serrano-Pérez-Higueras, F. Cavodeassi, S. Sotillos, M. D. Martín-Bermudo, A. Márquez, J. Buceta, L. M. Escudero, Scutoids are a geometrical solution to three-dimensional packing of epithelia. Nat. Commun. 9, 2960 (2018).

31. B. I. Shraiman, Mechanical feedback as a possible regulator of tissue growth. Proc. Natl. Acad. Sci. U. S. A. 102, 3318–3323 (2005).

32. S. Ithurbide, S. Gribaldo, S.-V. Albers, N. Pende, Spotlight on FtsZ-based cell division in Archaea. Trends Microbiol., doi: 10.1016/j.tim.2022.01.005 (2022).

33. O. Dudin, A. Ondracka, X. Grau-Bové, A. A. Haraldsen, A. Toyoda, H. Suga, J. Bråte, I. Ruiz-Trillo, A unicellular relative of animals generates a layer of polarized cells by actomyosin-dependent cellularization. Elife 8 (2019).

34. N. Fraikin, A. Couturier, C. Lesterlin, A palette of bright and photostable monomeric fluorescent proteins for bacterial time-lapse imaging, bioRxiv (2024) p. 2024.03.28.587235.

35. M. Olivetta, O. Dudin, The nuclear-to-cytoplasmic ratio drives cellularization in the close animal relative Sphaeroforma arctica. Curr. Biol. 33, 1597–1605.e3 (2023).

36. A. Ondracka, O. Dudin, I. Ruiz-Trillo, Decoupling of Nuclear Division Cycles and Cell Size during the Coenocytic Growth of the Ichthyosporean Sphaeroforma arctica. Curr. Biol. 28, 1964–1969.e2 (2018).

37. W. Seel, D. Baust, D. Sons, M. Albers, L. Etzbach, J. Fuss, A. Lipski, Carotenoids are used as regulators for membrane fluidity by Staphylococcus xylosus. Sci. Rep. 10, 330 (2020).

38. S. A. Sanchez, M. A. Tricerri, E. Gratton, Laurdan generalized polarization fluctuations measures membrane packing micro-heterogeneity in vivo. Proc. Natl. Acad. Sci. U. S. A. 109, 7314–7319 (2012).

39. H. Schiller, J. Kouassi, Y. Hong, T. Rados, J. Kwak, A. DiLucido, D. Safer, A. Marchfelder, F. Pfeiffer, A. Bisson-Filho, S. Schulze, M. Pohlschroder, Identification and characterization of structural and regulatory cell-shape determinants in Haloferax volcanii, bioRxiv (2023). 10.1101/2023.03.05.531186.

40. Z. Curtis, P. Escudeiro, J. Mallon, O. Leland, T. Rados, A. Dodge, K. Andre, J. Kwak, K. Yun, B. Isaac, M. Martinez Pastor, A. K. Schmid, M. Pohlschroder, V. Alva, A. Bisson, Halofilins as emerging bactofilin families of archaeal cell shape plasticity orchestrators. Proc. Natl. Acad. Sci. U. S. A. 121, e2401583121 (2024).

41. A. Charles-Orszag, N. A. Petek-Seoane, R. D. Mullins, Archaeal actins and the origin of a multi-functional cytoskeleton. J. Bacteriol. 206, e0034823 (2024).

42. S. Schulze, F. Pfeiffer, B. A. Garcia, M. Pohlschroder, Comprehensive glycoproteomics shines new light on the complexity and extent of glycosylation in archaea. PLoS Biol. 19, e3001277 (2021).

43. O. Vagin, J. A. Kraut, G. Sachs, Role of N-glycosylation in trafficking of apical membrane proteins in epithelia. Am. J. Physiol. Renal Physiol. 296, F459–69 (2009).

44. A. Liwosz, T. Lei, M. A. Kukuruzinska, N-glycosylation affects the molecular organization and stability of E-cadherin junctions. J. Biol. Chem. 281, 23138–23149 (2006).

45. T. Brunet, D. S. Booth, Cell polarity in the protist-to-animal transition. Curr. Top. Dev. Biol. 154, 1–36 (2023).

46. R. N. Esquivel, S. Schulze, R. Xu, M. Hippler, M. Pohlschroder, Identification of Haloferax volcanii pilin N-glycans with diverse roles in pilus biosynthesis, adhesion, and microcolony formation. J. Biol. Chem. 291, 10602–10614 (2016).

47. P. Lappalainen, T. Kotila, A. Jégou, G. Romet-Lemonne, Biochemical and mechanical regulation of actin dynamics. Nat. Rev. Mol. Cell Biol. 23, 836–852 (2022).

48. J. A. Theriot, Why are bacteria different from eukaryotes? BMC Biol. 11, 119 (2013).

49. P. K. Strother, M. D. Brasier, D. Wacey, L. Timpe, M. Saunders, C. H. Wellman, A possible billion-year-old holozoan with differentiated multicellularity. Curr. Biol. 31, 2658–2665.e2 (2021).

