## Supplemental Materials for "Tissue-Like Multicellular Development Triggered by Mechanical Compression in Archaea"

**The PDF file includes:**

Materials and Methods
Figs. S1 to S13
Tables S1 to S7
References

**Other Supplementary Materials for this manuscript include the following:**

Movies S1 to S17
Data S1 to S5

### Materials and Methods

Statistical Analysis

Experimental design, sample sizes, number of biological replicates, and statistical tests are described in Table S7. Briefly, distributions were represented as boxplots where the box limits define the interquartile range around the median; error bars represent a 95% confidence interval (CI). For XY graphs, shaded areas indicate the 95% CI. When appropriated, P-values and statistical significance was done using Kolmogorov-Smirnov (distributions with at least 50 datapoints without assumed normality) or One-Way ANOVA (comparisons between 2 groups with 3 datapoints each).

Haloarchaeal Strains

All haloarchaeal species (other than *Hfx. volcanii*, *Hfx. mediterranei*, *Hca. hispanica*, and *Hbt. salinarum*, which were kindly shared by Amy Schmid, Duke University) were provided by the DSMZ culture collection as described in Table S4.

Media and Culture Growth

All *Haloferax* species and strains were grown in the semi-defined Hv-Cab medium (51). Strains were streaked from -80°C freezer glycerol stocks onto Hv-Cab plates (1.5% agar), supplemented with 20 µM tryptophan and 50 µM uracil when necessary, and incubated for 2-3 days at 45°C. Single colonies were transferred into 16×25 mm glass tubes with 3 mL of liquid Hv-Cab, placed on a roller drum, and grown at 42°C to mid-exponential phase ($OD_{600nm}$~0.5). When necessary, the expression fluorescent reporters were induced by adding 500 µM Tryptophan. Other haloarchaeal species (as described in Table S4) were grown on 372-Cab medium, adapted from Hv-cab for other halophilic organisms: 3.3M NaCl, 56mM KCl, 180mM MgCl2, 174mM MgSO4, 10g/L Casaminoacids (Oxoid), pH 7.4, supplemented with thiamine 1 g/L, biotin 0.1 g/L, and trace elements (4.9mM $FeCl_3$, 0.37mM $ZnCl_2$, 0.074 mM $CuCl_2$, 0.077mM $CoCl_2$, 0.16mM $H3BO_3$, 12.7mM $MnCl_2$, 0.065mM $NiSO_4$, 0.041mM $Na_2MoO_4.2H_2O$).

Growth Curves

*Hfx. volcanii* strains were grown at 42°C until an $OD_{600nm}$=0.5, diluted 10-fold to an $OD_{600nm}$=0.05 and transferred to a 96-well flat-bottom plate (Corning Inc., #3370) next to Hv-Cab blanks. All wells surrounding the samples were filled with 200 µL $ddH_2O$ to prevent media evaporation. Growth curves of biological triplicates were performed using an EPOCH2 microplate spectrophotometer (Agilent) with constant orbital shaking at 42°C. $OD_{600nm}$ measurements were collected every 30 minutes for 48 hours. Each measurement was averaged across triplicates and then subtracted from the averaged Hv-Cab blank measurements.

Storage Modulus Measurements by Dynamic Mechanical Analysis (DMA)

Fresh pads under different agarose concentrations (0.25, 0.5, 1.0, 1.5, 2.0, 2.5, and 3.5%, w/v) were sent to Element Materials Technology (New Berlin WI) for Dynamic Mechanical Analysis (DMA). A small circle was cut from the provided agarose pad and placed within a Thermal

Analysis (TA) DMA Q800 instrument equipped with a compression clamp fixture. The Storage Moduli (E') were calculated using the equation (51):

$$E' = \frac{\sigma_0}{\gamma_0} \cos \delta$$

where $\sigma_0$ is stress, $\gamma_0$ is strain, and $\delta$ is the phase lag between stress and strain. The analysis was conducted at 1 Hz and an oscillating amplitude of 3 µm. Each sample was heated from 10°C to 80°C at a 2°C/min rate.

Membrane Fluidity Measurements with Laurdan

Cells were grown as described above and kept at the desired temperature (34$^{o}$C, 37$^{o}$C, or 42$^{o}$C) during the experiment unless indicated. To increase Laurdan permeability to *Hfx. volcanii* membranes, cells were spheroplasted by spinning down 1.5 mL of culture (2,500xg for 2 minutes) and resuspending the pellet in 100 µL of EBS Solution (1M NaCl, 25mM KCl, 0.4M Sucrose, 50mM EDTA, and 50mM Tris-HCl pH 8.5). After a 10-minute incubation at room temperature, 10% DMF and 100 µM Laurdan (Invitrogen, #D250) were added to spheroplasts, and tubes were mixed at maximum speed in a ThermoMixer Compact (Eppendorf) at desired temperatures (34$^{o}$C, 37$^{o}$C, or 42$^{o}$C) for 10 minutes. A 2 µL droplet of spheroplasted culture was transferred to a prewarmed 35mm glass-bottom dish (Ibidi, #81218-200) and gently covered with a pre-warmed 0.25% agarose pad prepared for S-layer staining as described above. Cells were then imaged on a Nikon Ti2E inverted microscope AX-R resonant scanning confocal with 405, 488, 561, and 640 nm lasers, aa CFI60x Apochromat oil objective and OKOlab caged incubator set to the desired temperature. Samples were imaged using the 640 nm laser (for S-layer) and the Spectral Detection for capture every 10nm from 420 nm to 670 nm (for Laurdan). For analysis, a binary mask was created for the S-layer image using the built-in Thresholding (Default setting). The mask was then applied to the individual Spectral Detection images after background subtraction, and the mean gray value was obtained for each cell.

*Hvo* and *Hmed* Viability under Micropillar Chambers

Mid-exponential *Hfx. volcanii* and *Hfx. mediterranei* liquid cultures ($OD_{600nm}$=0.5) were diluted 100-fold and 2 µl were transferred to 50 mm MatTek dishes (1.5mm coverslips) and compressed under microchambers fabricated as described in the "Micropillar Chambers" section. After confirming that cells were sparsely distributed (no more than one cell for every 10x10 µm pillar) by phase-contrast microscopy, samples were incubated at 42°C for 24 hours. Following incubation, cells were washed out from chamber with Hv-Cab liquid media, $OD_{600nm}$ adjusted to 0.1, and 100 µl from serial dilutions were plated onto Hv-Cab agar plates. After 48 hours, colonies were counted and colony unit formation (CFU) determined. Viability was plotted as values normalized by CFU from mid-exponential liquid cultures used as starting inoculum. For the viability comparison between *Hvo* colonies and tissues (Fig. S7D), 10 individual colonies were picked from Hv-Cab agar plate streaks, resuspended in liquid Hv-Cab, serial dilutions plated and CFU determined as described above.

Quantification of Biofilm Mass from *Haloferax* species

Biofilm was quantified with crystal violet as described in Schulze et al. (2022) with modifications. Briefly, biological triplicates of *Haloferax* species were grown in Hv-cab at 42°C to an $OD_{600}$~2.0, under constant agitation in a roller drum. Cultures were then diluted to an $OD_{600}$ of 1.5 and incubated at 42°C static for 60h. Cells were pelleted by centrifugation at 3000xg for 10 minutes and resuspended in 30 µL of Hv-cab (1 µL was used for serial dilutions, plated for CFU count). 200 µL of 2% acetic acid was added to the remainder of the cell suspension, which was then incubated at RT for 15 minutes, followed by centrifugation at 10,000xg for 2min. 200 µL of 0.1% crystal violet was added to the pellet, followed by vortexing to break the pellet, incubation at RT for 20min, and centrifugation at 10,000xg for 2min and two ddH2O washes. The pellets were dried for 20 minutes, followed by adding 500 µL of 30% acetic acid, and incubated at RT overnight, protected from light. 20 µL of the sample was then transferred to a 96-well plate, mixed with 180 µL of 30% acetic acid per well, and absorbance was measured using an EPOCH2 microplate spectrophotometer (Agilent) at 550nm. Absorbance values were normalized by colony formation units (CFU).

Quantification of Carotenoid Levels from *Hfx. volcanii and Hfx. mediterranei*

Carotenoid quantification was done as described in Cerletti et al (2014) with minor modifications. Briefly, cells were grown to stationary ($OD_{600}$ 1.5), centrifuged for 10 minutes at 4500xg and the pellet was resuspended in 1 mL of 1:1 Methanol:Chloroform and vortexed thoroughly for 10min. Samples were then centrifuged for 2min at 10000xg, and the 495nm and 475nm absorbance peaks of the supernatant were measured using a UV-Vis Spectrophotometer (Nanodrop,
ThermoScientific) and used to calculate carotenoid concentration.

RNA-seq of Multicellular Developmental Stages

DS2 cells were grown to an $OD_{600}$ of 0.15 in Hv-Cab and concentrated by centrifugation to an $OD_{600}$ of 0.6 in the same media. An aliquot of this cell culture was then saved for RNA extraction and corresponds to time 0h. Thirteen 3 µL droplets of cell culture were then placed equidistantly on glass Petri dishes (Corning 3160-101BO), and each was covered with a 1.5cm x 1.5 cm pad. The plates were washed with warm (42°C) Hv-cab for cell retrieval after 6h, 12h and 24h. Cells were immediately concentrated by centrifugation (4500xg for 10 minutes) and lysed with 1 mL of TriZol (Invitrogen) per sample. Total RNA extraction was performed as described (Rados and Andre, 2023). Samples were sent to SeqCenter LLC (Pittsburgh, USA, seqcenter.com) for ribosome depletion using *Haloferax*-specific probes (Rados and Andre, 2023) and sequencing. Pre-processing and quality filter of paired-end raw sequencing reads were performed by fastp (v.0.23.4) (53) under default arguments. Reads that passed this quality filtering step were eligible for downstream transcript quantification using Salmon (v.1.10.0) (54) in a two-step procedure.

First, the nucleotide CDS sequences for the *Haloferax volcanii* genome ("_cds_from_genomic.fna.gz" suffix file) were downloaded from the NCBI Genomes FTP website (http://ftp.ncbi.nlm.nih.gov/genomes/all/), using the respective assembly and RefSeq accessions
(Table S1). To build an index of the *Haloferax volcanii* transcriptome, the "salmon index" command was run using the file above as input and default arguments. Second, transcripts were quantified with the "salmon quant" command, using the quality-filtered reads and the index built in the previous step as input. The sequence-specific bias correction parameter was enabled, and the library type was set to be automatically inferred. All other parameters were left to default. A gene count matrix with dimensions "number of samples" versus "number of genes" was created with the read counts generated by Salmon. Genes with less than 10 reads across all samples were discarded from this matrix.

Differential Gene Expression (DGE) analysis

DGE analysis was performed with PyDESeq2 (v.0.4.9) (55), using the gene count matrix as input, together with metadata concerning the experimental design. Gene expressions were compared between the reference (unicells, T=0h) and the samples collected at the 6h (coenocytes), 12h (cellularization), and 24h (tissues) time points. Genes whose $|\log2(\text{Fold Change})| \geq 1$ and adjusted p-value $\leq 0.05$ were used for downstream data analysis and visualization. Data analysis was carried out with Pandas v.1.5.3 (56), and plots were created using Matplotlib v.3.7.1 (57). These packages were installed and run under Python v.3.10.12 (58).

Tree of *Haloferax* Species

Annotated protein sequences from Halobacteria genomes were retrieved from the NCBI GenBank database, out of the 315 genomes 78 are part of the *Haloferax* genus. The accession numbers and associated metadata for these genomes are provided in Supplementary Data S5 and Data S6.

Homologous gene clusters were identified using GET_HOMOLOGS's (59) implementation of the OrthoMCL algorithm (60). A total of 385 core homologous groups are present in at least 90% of the studied Halobacteria genomes. Each homologous group was independently aligned using MAFFT (61) with --reorder and --auto parameters. Once aligned, sequences were concatenated into a single supermatrix comprising 162,041 sites.

A phylogenetic tree was reconstructed using IQ-TREE (62), and the substitution model LG+F+I+G4. Bipartition support was estimated using both the Ultrafast Bootstrap (i.e., UFBoot) with 1,000 replicates (63) and the SH-like approximate likelihood-ratio test (SH-aLRT) with 1,000 replicates (64). The unrooted maximum likelihood tree generated by IQ-TREE was rooted using the Minimal Ancestor Deviation (MAD) method (65) to objectively determine the root position within Halobacteria without the need for an outgroup. MAD version 2.0 was employed with default settings, minimizing the deviation in root-to-tip lengths across the tree. Software and data are available in the following suppositories: GET_HOMOLOGUES (66), MAFFT (67), Q-TREE (68), and MAD (69).

Comparative Genomics Analysis of *Haloferax* Species

The protein sequences ("_protein.faa.gz" suffix file) for each of the 11 *Haloferax* species' genomes shown in Table S1 were downloaded from the NCBI Genomes FTP website (70), using the assembly and RefSeq accessions respective to each species. OrthoFinder (v.2.5.5) (71) was run using the protein sequences from these 11 genomes as input, with default arguments. OrthoFinder was also run separately for two other datasets: (i) one containing the protein sequences from the *Haloferax volcanii*, *Haloferax gibbonsii*, *Haloferax prahovense*, and *Haloferax mediterranei* genomes, and (ii) another containing the protein sequences from the *Haloferax volcanii*, and *Haloferax mediterranei* genomes. Genomic features for the *Haloferax volcanii* protein sequences within each set of orthologous sequences (orthogroup) were gathered from the "_feature_table.txt.gz" file for the respective genome (see Table S1). This file was downloaded from the NCBI Genomes FTP website. The "idmapping.dat" file was downloaded from the UniProt FTP website (72). Next, the RefSeq id of each protein sequence was cross-referenced against it to retrieve the respective UniProt id. Data analysis was carried out on the output files generated by OrthoFinder using Pandas v.1.5.3 (56), installed and run under Python v.3.10.12 (58).

Over-Representation Analysis (ORA) of Orthologous Protein Groups

The protein counts generated by OrthoFinder were used for all orthogroups and each of the 11 Haloferax species in Table S1. Our dataset of 11 species was divided into two groups: (i) species that form tissue and (ii) species that do not form tissue (Table S1). The null hypothesis H0 was defined as orthogroup membership and belonging to the group of species that form tissue are independent properties, and the alternative hypothesis H1 as there is an over-representation of proteins from an orthogroup that belong to the group of Haloferax species that form tissue. This problem was formulated as a contingency table (see Table S2). The test performed was the one-sided hypergeometric test, and the critical region chosen was the upper tail of the distribution. To calculate the p-value associated with this test and for each orthogroup, the survival function of the hypergeometric distribution was used. To control the false positive rate (FDR) from these multiple comparisons, the Benjamini-Hochberg correction (73) was applied to all p-values calculated this way. The over-represented orthogroups that satisfied all the following conditions were considered for downstream data analysis and interpretation: (i) the orthogroup has at least one protein from the Haloferax volcanii genome; (ii) the orthogroup has proteins from at least 4 *Haloferax* species that form tissue; (iii) the orthogroup has a hypergeometric test FDR-adjusted p-value $\leq 0.05$. Data analysis was carried out with Pandas v.1.5.3, hypergeometric tests were performed with SciPy
v.1.14.0, and the Benjamini-Hochberg correction to the p-values was achieved with statsmodels v.0.14.2. These packages were installed and ran under Python v.3.10.12.

Cryo-EM and cryo-ET of cells from sheared tissues

Grids for cryo-EM experiments were prepared as described previously (74). Briefly, 2.5 μL of *Hfx. volcanii* unicells were supplemented 10:1 (cells:gold solution, v/v) with 10 nm protein-A gold (CMC Utrecht) immediately prior to grid preparation. For sheared multicells, *Hfx. volcanii* cells were grown under compression for 16 to 17 hours, then washed with the Hv-cab media supplemented 8:1 (cells:gold solution, v/v) with 10 nm protein-A gold (CMC Utrecht). These mixtures were then applied to a freshly glow discharged Quantifoil R2/2 (washed multicells) or R3.5/1 (unicells) Cu/Rh 200 mesh grid, adsorbed for 10 s, blotted for 4-5 s and plunge-frozen into liquid ethane that was maintained at -178 ºC in a Vitrobot Mark IV (ThermoFisher), while the blotting chamber was maintained at 100% humidity at 10 ºC. The frozen grids were stored in liquid nitrogen until further investigation.

Cryo-ET data was collected as described previously (75-77). Briefly, for tomographic data collection, the SerialEM software (78) was used in a Titan Krios microscope equipped with a Quantum energy filter (slit width 20 eV) and K3 direct electron detector running in counting mode.

Tilt series with a defocus range of −3 to −6 μm (washed multicells) or -5 to -8 μm (unicells) were collected between ±60° in a dose symmetric scheme (79) with a 3° (washed multicells) or 2° (unicells) tilt increment. A total dose of 80 e-/Å2 (washed multicells) or 85 e-/Å2 (unicells) was applied over the entire series. Cryo-ET data analysis: tilt-series alignment and tomographic reconstruction were carried out using IMOD (80). Datasets were motion-corrected and doseweighted with MotionCor2 implemented in Relion 5.0 (81). Contrast transfer functions (CTFs) of the resulting motion-corrected micrographs were estimated using CTFFIND4 (82). Data visualization was performed in IMOD.

Live-Cell Microscopy and Image Analysis

**Immobilization and Compression of *Hfx. volcanii*:** Agarose pads were prepared by mixing Hv-Cab and Seakem agarose (Seakem Lonza Inc 50002) and autoclaving bottles in a pressure cooker (Instant Pot Duo 8-quart V5) for 30 minutes. Homogenous agarose solutions were immediately poured into 100x15mm Petri dishes (Corning™ FisherSci 07-202-030) as a mold and left to polymerize at room temperature overnight. Plates were wrapped in parafilm and stored at 4°C inside Ziploc bags. For tissue development, colonies from fresh plates (under one week old) were grown in liquid until $OD_{600nm}$~0.2. 3 μL culture droplets were placed onto clean 50 mm Mattek dishes (MatTek Corp., #P50G-1.5-30-F), gently covered with a (unless otherwise indicated) 1.5cm x 1.5 cm 2.5% agarose pad and incubated at temperatures and times indicated in each experiment.

**S-layer and ConA-Alexa488 Staining:** S-layer was stained with Brilliant Blue FCF, a nontoxic triarylmethane dye impermeable to cells (83) that binds preferentially to glycoproteins. Brilliant Blue FCF was added at a 1:4000 concentration to liquid cultures (Brilliant Blue FCF,

Sigma, 3844-45-9) or directly to pads (Brilliant Blue dye, Spice Supreme, NJ) after autoclaving. For ConA-Alexa488 staining, 30 µL of ConA-Alexa 488 1 mg/mL conjugate stock (Invitrogen, C11252) was added to the top of the pad and allowed to diffuse for 8 hours before imaging.

**Coverslip Cleaning for Super-Resolution and Single-Molecule Imaging:** Coverslips were placed in a custom 3D-printed tray, sonicated for 15 minutes in 2M KOH, and rinsed with MilliQ water. They were sonicated again for 15 minutes in 4% Extran MA 02 detergent (Millipore Sigma, #1075532500) and rinsed with Milli-Q water. They were then sonicated a third time for 15 minutes in 190-proof, benzene-free ethanol, rinsed with ethanol, and parafilm-wrapped and stored in fresh ethanol. Before imaging, coverslips were fished from the container with clean tweezers, sonicated for 5 minutes in ethanol, and dried with a compressed air duster.

**Image Segmentation:** For time-lapse experiments requiring lineage tracking (ArcCell device, cellularization control, PCNA foci counting, etc), images were segmented using Omnipose (84) with a custom in-house trained model. Segmented images were converted to binary masks, and CellProfiler (85) was used to measure and track cells over time. Unicells with areas above 10 $\mu m^2$ or circularity below 0.7 were filtered out for quality control.

**Epifluorescence and Phase-Contrast Microscopy:** Cells were imaged using a Nikon TiE2 inverted microscope equipped with a Lumencor Sola II Fluorescent LED (380-760 nm), Hamamatsu ORCA Flash 4.0 v3 sCMOS Camera (6.5 µm/pixel), and a CFI PlanApo Lambda 100x DM Ph3 oil objective (NA=1.45). The microscope body was enclosed by an OkoLab Caged incubator set to temperatures indicated in each experiment.

**SoRa Microscopy:** Cells were grown as described, and a 3 µL droplet was placed on a 35mm glass-bottom dish (Ibidi, #81218-200) and gently covered with a pre-warmed 2.5% agarose pad prepared for S-layer staining as described above. Samples were then incubated at 42$^o$C for 6h (coenocytes), 10-12h (cellularization), or 18-20h (tissues). Super-resolution images were collected at room temperature using a Nikon Ti-2 equipped with a Yokogawa CSU-W1 SoRa spinning disk, with lasers 640 nm (Brilliant Blue) and 488 nm (GFP), a Plan Apo λ 100x objective (NA=1.45), and a Prime BSI express camera (6.5 µm/pixel), with a Z-step of 0.1µm. Images were processed using NIS-Elements software (Nikon), using NIS.ai for Richardson-Lucy 3D deconvolution.

**iSIM microscopy:** Cells expressing cytoplasmic GFP were grown as described, and a 3 µL droplet was placed on a 35mm glass-bottom dish (Ibidi, #81218-200) and gently covered with a pre-warmed 2.5% agarose pad prepared. Samples were then incubated at 42$^o$C for 18h. Images were collected at room temperature using a Leica microscope equipped with an iSIM 488 Quad, a

Plan Apo λ 100x objective (NA=1.45) with a Z-step of 0.2µm. Images were processed using NISElements software (Nikon), using NIS.ai for Richardson-Lucy 3D deconvolution.

**STED Microscopy:** *Haloferax volcanii* WT cells (DS2) were grown as described. A 3 µL droplet was placed on a 35mm glass-bottom dish (Ibidi, #81218-200) and gently covered with a pre-warmed 2.5% agarose pad prepared for S-layer staining as described above. Samples were then incubated at 42$^{o}$C for 14-16h. Super-resolution images were captured at room temperature on an Abberior 3D STED system, mounted on an Olympus IX83 microscope with 405, 485, 561, and 640 nm excitation lasers and a 60x oil immersion objective. Images were segmented using a custom in-house pre-trained Cellpose (86-87) model available at https://github.com/Archaea-Lab/Multicell_paper (88). Volume and surface area were calculated using Fiji (89). Segmentation outlines were processed and rendered using Imaris 10.2 (Oxford Instruments).

**Traction Force Microscopy (TFM):** Fluorescent polystyrene beads of 50 nm diameter (Thermo Scientific, #09-980-410) were diluted 1:100 in a 0.0125% Tween 20 (Thermo Fisher #28320) solution and vortexed. Petri dishes (Genessee, 32-105G) were plasma treated for two minutes. A 200 µL aliquot of the bead solution was pipetted directly on the Petri dish and allowed to bond for 15 seconds. The Petri dish was inverted to pour off excess solution and then allowed to air dry. *Hfx. volcanii* cells were grown to an $O.D._{600nm}$ of 0.14-0.20, and 1 µL of media was pipetted beneath a bead-embedded 2.5% agarose pad, bead side down. 3D-stack timelapses were collected at 1-hour intervals by both phase-contrast and epifluorescence. 3D-stacks were collected with 34 slices at 0.2 µm intervals.

**Recompression of Cells from Fragmented Tissues:** Hvo tissues under 2.5% agarose pads were placed under the microscope and imaged by phase-contrast microscopy. With ongoing timelapses, tissues were broken by shear-pipetting of 50 µl of Hv-Cab media in between pads and coverslips. After having cells ruptured from tissues, cells were recompressed by removal of excess liquid from pad using a kimwipe paper tissue touching the top of the pad around the imaged area. Cells were segmented and tracked as described above, and deformation was calculated as the difference recompressed cell area ($A_k$) and initial uncompressed cell area ($A_0$), normalized by the initial cell area ($A_0$). Annotation of peribasal and apicobasal cells was done manually across tissues.

**Volactin Filament Angle Analysis:** aBL126 cells were grown as described, and a 3 µL droplet was placed on a 35mm glass-bottom dish (Ibidi, #81218-200). For tissue analysis, cells were gently covered with a pre-warmed 2.5% agarose pad prepared for S-layer staining, as described above. For single-cell analysis, the process was repeated with a 0.5% agarose pad. Samples were then immediately imaged (single cells) or incubated at 42$^{o}$C for 6h (coenocytes), 10-12h (cellularization), or 18-20h (mature tissues). Images were collected at room temperature using a Nikon Ti-2 equipped with a Yokogawa CSU-W1 SoRa spinning disk, with lasers 640 nm (Brilliant Blue) and 488 nm (GFP), a Plan Apo λ 100x objective, and a Prime BSI express camera (6.5 µm/pixel), with a Z-step of 0.1µm. Filament angles were calculated manually using Fiji straight-line tool.

**Laser Ablation:** Cells were imaged at room temperature using a Nikon Ti-2 equipped with a Yokogawa CSU-W1 spinning disk, with lasers 640 nm (Brilliant Blue), with a Plan Apo λ 100x objective, Prime BSI express camera (6.5 µm/pixel). Different ROIs were selected, and 100mslong ablation pulses were performed using a 405 nm laser. Cells were then subsequently imaged under 30-second intervals for 15 minutes (large area ROIs) or 3-second intervals for 15 seconds (small area ROIs). Mean Squared Displacement (MSD) was calculated by processing timelapse images segmented with a pre-trained Cellpose model (86-87). The resulting image stacks were analyzed using the Fiji plugin TrackMate (Ershov et al, 2022), with the built-in Label Image Detector and Simple LAP Tracker. The resultant masks were exported as ImageJ ROIs to determine the center of mass (COM) for each independent cell. The initial ablation coordinates, along with the COM coordinates from the first and last frames of each cell, were then used to calculate the MSD for each cell. For plotting and analysis, only cells that were moving directionally ($\alpha>1$) and towards the ablation point ($\Delta$distance<1) were considered. The custom Python script for MSD calculations and the pre-trained Cellpose models used can be found at https://github.com/Archaea-Lab/Multicell_paper (88). This approach was modified from a thread in the imageJ.sc forum (90).

**Microfluidics:** The ArcCell chips are produced using standard photolithographic techniques (91). The silicon wafer was coated by a negative tone photoresist (SU-8). After soft backing, the desired pattern was exposed on the wafer via direct writing. The devices have channels with distinguished lengths, widths, and heights (Fig. S1A). After the master was ready, it was cast with polydimethylsiloxane (PDMS) degas and cured at 70°C. When cured, the PDMS was cut, peeled and punched to connect ports to the external tubing. The ready mold was then bonded irreversibly to cover glass using oxygen plasma. The ArcCell device was designed to ensure high cell loadings in the chamber and low shear prefusion of fresh media to cell trapping areas. The device has two working modes: loading and running. The loading output is open, the run output is blocked, and cells and media are gently forced to flow into the trapping chambers. After loading, the run output is open, and the media flow rate at the chamber is reduced. Loading and flow rates in the ArcCell device were performed using the LU-FEZ-0345 FlowEZ Microfluidic Flow Controller (Fluigent) coupled with a 0-1 mL/min Flow Unit L (Fluigent). Cells were loaded under 1 µL/min flow rates, and time-lapses were recorded under a constant flow rate of 2 µL/min.

**Micropillar Chambers:** The squared microchambers with micropillars were designed using the open software tool KLayout with planar dimensions of $10\times10\mu m^2$. The microfabrication process was previously described in (92-94). Briefly, 3” silicon dioxide wafers (Siegert Wafer GmbH, Germany) were extensively cleaned in a solution of 1:1:5 $H_2O_2$:$NH_4OH$:$H_2O$ at 70°C for 10 min before use. After drying, the clean wafers were incubated with a 1:10 hexamethyldisiloxane:isopropanol priming solution directly on top. The priming solution was then removed by spincoating and the wafer was heated on a hot plate to 95°C for 5min on a heating plate to remove excess solvent. The hexamethyldisiloxane primer is activated at 95°C for a further 5min to improve adhesion of the positive photoresist Shipley 1813 (Micro Resist Technology GmbH, Germany). The photoresist is spincoated on the silicon substrate to achieve a height of 1.3 µm according to the manufacturer instructions. After curing, the squared microchamber design was written directly onto the photoresist using a tabletop µMLA system (Heidelberg Instruments, Germany). After exposure, the photoresist was developed in MF-321 (Micro Resist Technology GmbH, Germany) for ~2 min with gentle shaking and dried under a stream of nitrogen. The resulting master template was silanized overnight under vacuum with a drop of heptadecafluoro1,1,2,2-tetrahydrodecyl trichlorosilane (abcr GmbH, Germany). The master design was then replicated into PDMS (Sylgard184) using a ratio of 10:1 base:curing agent, degassed and cured overnight at 65°C under vacuum. Subsequently, the PDMS mold was then separated from the silicon master and replicated in a growth medium-agarose mixture for growth experiments.

***Hvo* and *Hmed* Viability under Micropillar Chambers:** Mid-exponential *Hfx. volcanii* and *Hfx. mediterranei* liquid cultures ($OD_{600nm}$=0.5) were diluted 100-fold and 2 µl were transferred to 50 mm MatTek dishes (1.5mm coverslips) and compressed under microchambers fabricated as described above. After confirming that cells were sparsely distributed (no more than one cell for every 100x100 µm area) by phase-contrast microscopy, samples were incubated at 42°C for 24 hours. Following incubation, cells were washed out from chamber with Hv-Cab liquid media, $OD_{600nm}$ adjusted to 0.1, and 100 µl from serial dilutions were plated onto Hv-Cab agar plates. After 48 hours, colonies were counted and colony unit formation (CFU) determined. Viability was plotted as values normalized by CFU from mid-exponential liquid cultures used as starting inoculum. For the viability comparison between *Hvo* colonies and tissues, 10 individual colonies were picked from Hv-Cab agar plate streaks, resuspended in liquid Hv-Cab, serial dilutions plated and CFU determined as described above.

**Growth Rate and Lifespan Measurements:** Epifluorescence and phase-contrast timelapses (20-minute intervals) of *Hvo* S-Layer-stained unicells were 3D- and 2D-segmented using CellPose2 (86) model available at https://github.com/Archaea-Lab/Multicell_paper (88). Segmented images were converted to binary masks, and CellProfiler (85) was used to create lineages tracks to extract single-cell volume and surface area over time. Each lineage was checked manually and traced back to their progenitor cells. The lifespan of each cell was defined as the time between its origin and its next division. Cells that did not complete their entire lifespan within the timelapse were excluded from the analysis. Growth rate was then calculated by dividing the area added by lifespan.

**bSpoJ Particle Tracking:** bSPoJ expression was induced under 200uM tryptophan and grown to exponential phase ($O.D._{600nm}$~0.2). For bSpoJ tracking in unicells, cultures were incubated with 75-150pM HaloTag-ligand JFx549 (kindly provided by Luke Lavis and lab at

Janelia) for 30 minutes under standard growth conditions. The cultures were concentrated 10-fold (3,000 xg for 3 min), and 2 µL droplets of culture were placed on 60×24 mm sonicated-clean coverslips and gently immobilized under pre-warmed 1.5x0.5 cm, round 0.5% agarose pad. For bSpoJ tracking in early and cellularizing coenocytes, cells from liquid exponential cultures were compressed under 2.5% agarose pads prepared with 200 µM tryptophan and 50-500pM JFx549, placed onto a clean 50 mm Mattek dish (MatTek Corp., #P50G-1.5-30-F) and incubated at 42°C for ~6 (early coenocytic phase) and ~12 (onset of cellularization) hours protected from light. TIRF time-lapses were collected in a Nikon TiE2 inverted microscope equipped with a Hamamatsu ORCA Flash 4.0 v3 sCMOS Camera (6.5 µm/pixel), a CFI PlanApo Lambda 100x DM Ph3 oil objective. The microscope body was enclosed by an OkoLab Caged incubator set at 42°C. TIRF time-lapses were collected under 500ms (unicells) or 50ms (coenocytes) exposure times and 2% laser power from a 70 mW Nikon LUN-F 561 nm laser line.

**bSpoJ Particle-Tracking Analysis:** Phase-contrast snapshots were automatedly segmented using Omnipose (84) using previously custom-trained model (44). Cell masks were exported from Omnipose and merged with the fluorescence TIRF channel. Foci positions were tracked, and diffusion speed was calculated in Fiji using the TrackMate plugin (95). Spot detection was performed with the LoG detector and the following parameters: Object Diameter=0.45 µm, Quality Threshold=0.2, Pre-process with median filter checked, and sub-pixel localization checked. Spots were filtered based on minimum intensity to exclude spots outside of cells. Tracking was performed with the Simple LAP Tracker and the following parameters: Linking Max Distance=1 µm, Gap-closing max distance=0 µm, Gap-closing max frame gap=0. Spot files, containing position information over time, and Track files, containing mean speeds of tracks (µm/s), were exported from TrackMate. Next, tracks from TrackMate were used to calculate Mean Squared Displacements (MSD). First, tracks with a lifetime below 5 frames were filtered out to preserve temporal information. The time-averaged MSD for a given time lag was calculated for each individual track following the equation:

$$\langle MSD(\tau)\rangle = \frac{1}{N}\sum_{t=t_{\text{start}}}^{T-\tau}\left[(x(t+\tau)-x(t))^2 + (y(t+\tau)-y(t))^2\right]$$

where x and y are the positions in 2 dimensions (µm), τ denotes the time lag interval between each position, t denotes the current time point of the track, $t_{start}$ is the first-time interval of the track, T is the total time in the track, and N-1 is the number of positions used in the current MSD calculation. For an individual track, the MSD was calculated with increasing time lags starting with the frame rate at which the movie was collected and ending with the total lifetime of the track. The effective diffusion coefficient, $D_{eff}$, was subsequently extracted from the linear fit to the truncated TA-MSD containing the first 5-time lags, following the equation:

$$MSD(\tau) = 4D_{eff}\tau$$

Tracks with fits $R^2<0.9$ were discarded. All calculations were done using a custom Python script (88).

Cloning and Transformation

Oligos, plasmids, and strains used in this study can be found in Tables S3-S5. Plasmids were cloned using Gibson assembly (96) and transformed into competent *E. coli* DH5α cells, transformants selected on LB plates supplemented with 100 µg/mL Carbenicillin. Plasmids were transformed into *Hfx. volcanii* using the method previously described (97). Unless noted, *Hfx. volcanii* strains were built from the parental H26 (Δ*pyrE2*) or DS2 wild-type strains. A detailed description of each strain created in this work follows:

*aSB14 [pcna::mevR-HaloTag-mNeonGreen-PCNA]* was created by transforming the DS2 strain with the eSB14 plasmid, cloned from a Gibson assembly consisting of four fragments: 1) pcna upstream region: PCR with primers oSB33 and oSB42 from DS2 genomic DNA template; 2) mevR-Ppcna: synthetic fragment carrying the mevR resistance cassette and 500bp sequence from the pcna promoter region; 3) HaloTag-15aa-mNeonGreen-pcna: synthetic fragment carrying both HaloTag and mNeonGreen tags and the first 750bp of the pcna gene; 4) pTA131 plasmid linearized with NotI. DS2 transformants were selected on Hv-Cab plates supplemented with 10 µg/ml mevinolin and uracil.

*aBL582 [ΔpyrE2 ftsZ1::ftsZ1-mChartreuse]* was created by transforming H26 with the eBL416 suicide plasmid, cloned from a Gibson assembly consisting of 4 fragments: 1) ftsZ1 fragment: PCR with primers oJM193 and oAB130 from DS2 genomic DNA template; 2) 40aamChartreuse synthetic fragment; 3) ftsZ1 downstream fragment: PCR with primers oBL532 and oBL533 from DS2 genomic DNA template; 4) pTSD1 plasmid, carrying the pyrE2 expression cassette, linearized with EcoRI. The pTSD1 suicide vector was created from a Gibson assembly consisting of a PCR fragment with primers oBL536 and oBL537 from the pTA131 vector, inserting an EcoRI restriction site upstream to the pyrE2 cassette. H26 cells were transformed, plated on Hv-Cab plates, and incubated at 45°C. Transformants were grown in liquid Hv-Cab overnight, back-diluted into Hv-Cab, and grown overnight. Cultures were then back diluted into Hv-Cab supplemented with 50 µM uracil, grown overnight, and then back diluted again in HvCab supplemented with 50 µM uracil. Cultures were struck onto Hv-Cab plates supplemented with 50 µM uracil and 50 µg/mL 5-FOA. Colonies were confirmed by PCR for the presence of *ftsZ1mChartreuse* and the absence of untagged *ftsZ1*.

*aOL39 [ΔpyrE2 pTA962::sec-spoIIIJ-HaloTag]* was created by transforming the H26 strain with the eOL52 plasmid, cloned from a Gibson assembly consisting of four parts: 1) SEC fragment: PCR with primers oHV13 and oOL79 from a synthetic DNA template; 2) spoIIIJ fragment: PCR with primers oOL80 and oOL81 from a PY79 (*Bacillus subtilis*) gDNA template; 3) 30aaHaloTag: PCR with primers oBL354 and oBL318 from a synthetic DNA template; 4) pTA962 plasmid linearized with NdeI. H26 cells were transformed in Hv-Cab plates.

*ID112 [ΔhdrB ΔpyrE2 ΔftsZ1 ΔftsZ2]* was a gift from Iain Duggin (UT Sydney).

ID77 *[ΔhdrB ΔpyrE2 ΔftsZ1]* was a gift from Iain Duggin (UT Sydney).

*Δ2528 [ΔpyrE2 ΔHVO_2528]* was a gift from Jörg Soppa (Goethe-Universität).

Δ*aglB [ΔpyrE2 ΔtrpA tn::aglB]* was a gift from Jerry Eichler (Ben Gurion University)

Δ*aglE [ΔpyrE2 ΔtrpA tn::aglE]* was a gift from Jerry Eichler (Ben Gurion University)

Δ*aglJ [ΔpyrE2 ΔtrpA tn::aglJ]* was a gift from Jerry Eichler (Ben Gurion University)

Δ*aglG [ΔpyrE2 ΔtrpA tn::aglG]* was a gift from Jerry Eichler (Ben Gurion University)

Δ*aglI [ΔpyrE2 ΔtrpA tn::aglI]* was a gift from Jerry Eichler (Ben Gurion University)

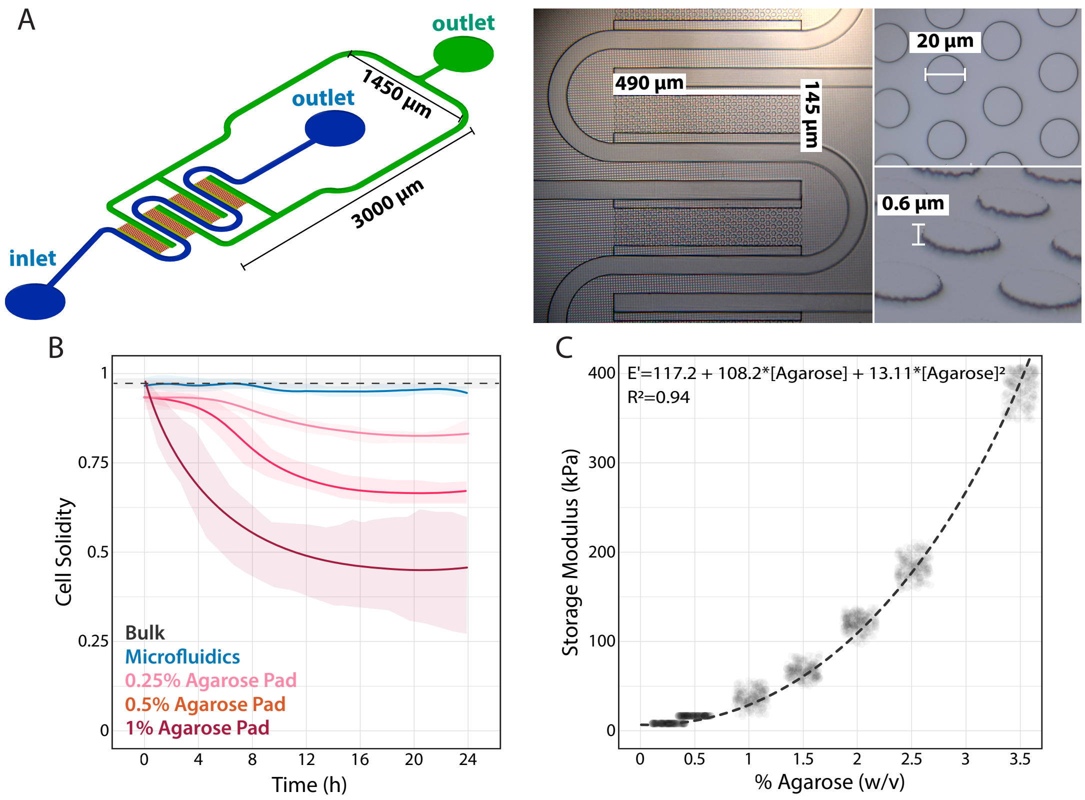


**Fig. S1. Different setups used for *Haloferax volcanii* live-cell imaging (A**) Schematic illustration of the ArcCell microfluidic device. Cells are trapped in the orange region. Media flows through the blue channels, diffusing into the orange region. The green region is an alternative outlet, allowing for the flow path to be manipulated into loading cells into the traps (left). Dimensions of the cell trapping region (middle). Dimensions of the pillars that create the height of the cell trapping region (right). Dimensions of the device were measured using 200 steps on a profilometer. **(B)** Cell solidity measurements of cells from liquid cultures (black), trapped in ArcCell (blue) or confined beneath 0.25%, 0.5%, and 1% agarose pads (pink gradient). **(C)** Storage moduli (E') of agarose pads measured by DMA. Data was fit under a second-order polynomial curve.

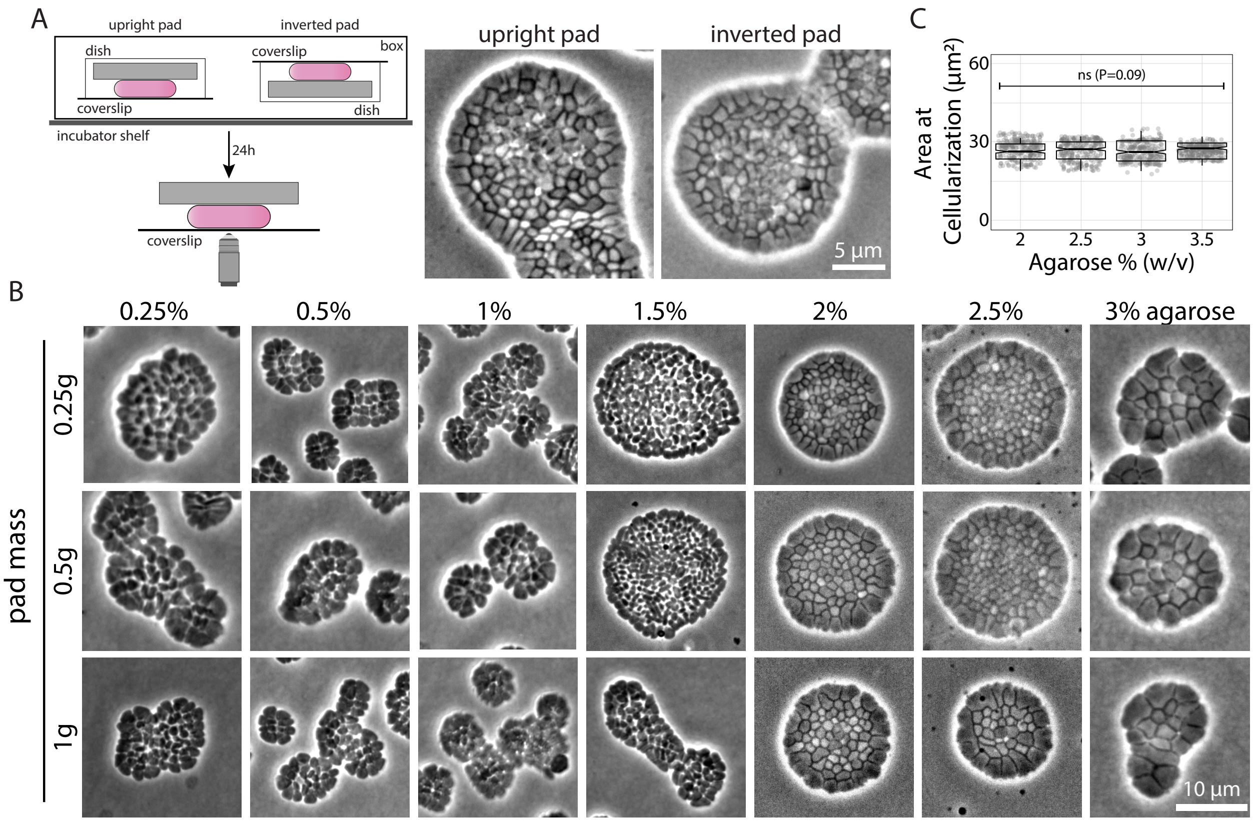


**Fig. S2. Tissue development is a precisely controlled process independent of pad weight, mass, and thickness. (A)** (left) Illustration of *Hvo* cells compressed by pads in upright and upside-down positions. (right) Phase-contrast images of tissues after 24 hours incubation at 42°C. **(B)** *Hvo* cells compressed under pads with different agarose concentrations and masses. Pad thickness scaled with masses as their areas were kept the same. **(C)** Cell area at cellularization is independent of agarose concentration above the 2% threshold.

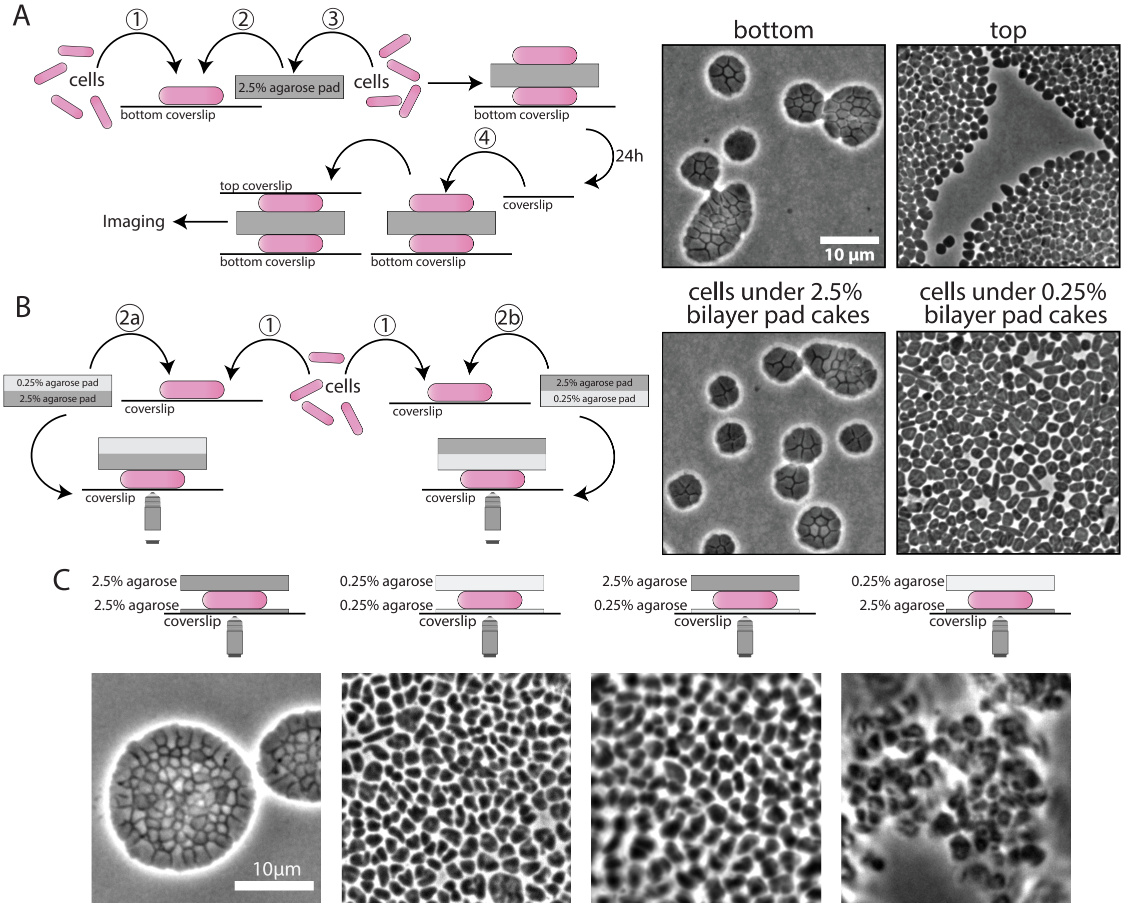


**Fig. S3. Pad stiffness is not sufficient to trigger tissue development (A-C)** Schematic illustrating the experiment performed to decouple substrate rigidity and compression. **(A)** (Left) *Hvo* cells were placed on a coverslip (step 1) and confined with a 2.5% agarose pad (step 2). The same culture was placed atop the agarose pad (step 3) and allowed to develop for 24 hours, unconfined. A secondary coverslip was placed on top of the agarose pad (step 4). (Right) Phase contrast images of cells on the bottom or top of the pad. **(B)** (Left) Bilayer agarose pads were poured with 2.5% on the bottom and 0.25% agarose on top. *Hvo* cells were then placed on two separated coverslips (step 1). The cells were compressed with either the 2.5% (step 2a) or 0.25% (step 2b) agarose surface facing towards them. (Right) Phase contrast images of cells under the 2.5% or 0.25% pad surface of the bilayer pad cakes. **(C)** *Hvo* cells were compressed in between an agarose pad (top) and a thin layer of agarose film coating the coverslip (bottom).

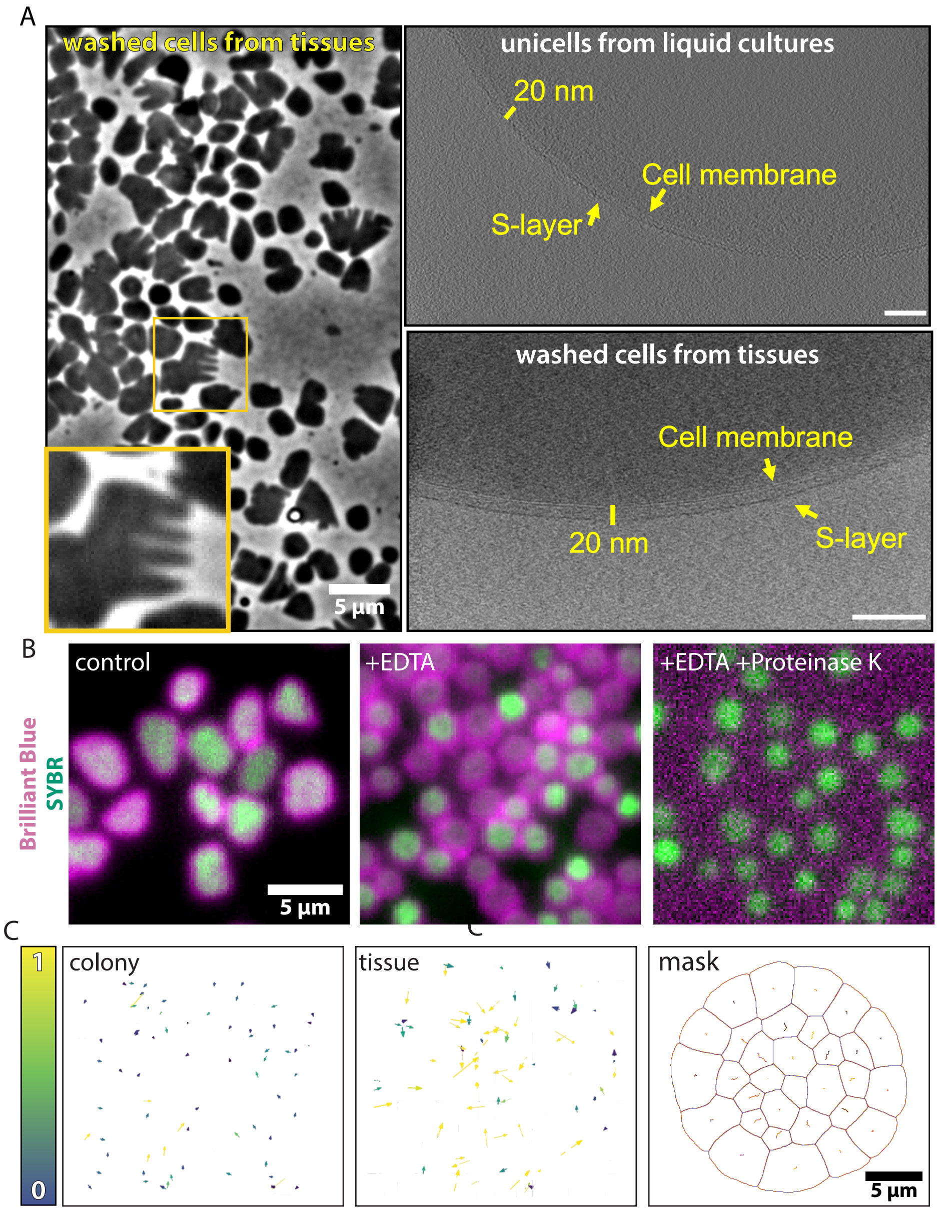


**Figure S4. Tissues maintain intact S-layer lattices but show distinct viscoelastic interaction properties. (A)** Phase-contrast micrographs of *Hvo* cells fragmented from tissues (left) and slices through a tomogram of a *Hvo* unicell (top-right) and a *Hvo* cell from, fragmented tissues (bottom-right), highlighting the cell membrane, S-layer and the distance measured between them, which agrees with previously published data (~12 nm distance from the membrane to the S-layer base and a further 8-10 nm thickness of the S-layer) (74). Scale bars: 100 nm. **(B)** Epifluorescence microscopy of *Hvo* unicells stained with Brilliant Blue (S-layer, magenta) and SYBR (DNA, green). (Left) Control cells from cultures in Hv-Cab medium. (Middle) Spheroplasting under 50 mM EDTA (10-minute incubation) disrupts the Slayer lattice but does not detach the S-layer glycoprotein from the cytoplasmic membrane. Cells under EDTA treatment show high heterogeneity in DNA staining. (Right) Spheroplasting under 50 mM EDTA and 1 mg/ml Proteinase K degrades outer-surface proteins and prevents Brilliant Blue staining. **(C)** (Left) Displacement vectors of cells' center of mass in response to laser ablation in the center of unicells (left) and a tissue (middle). (Right) Segmented cell masks and displacement tracks in a representative tissue. False-colored vectors and tracks were normalized from MSD values.

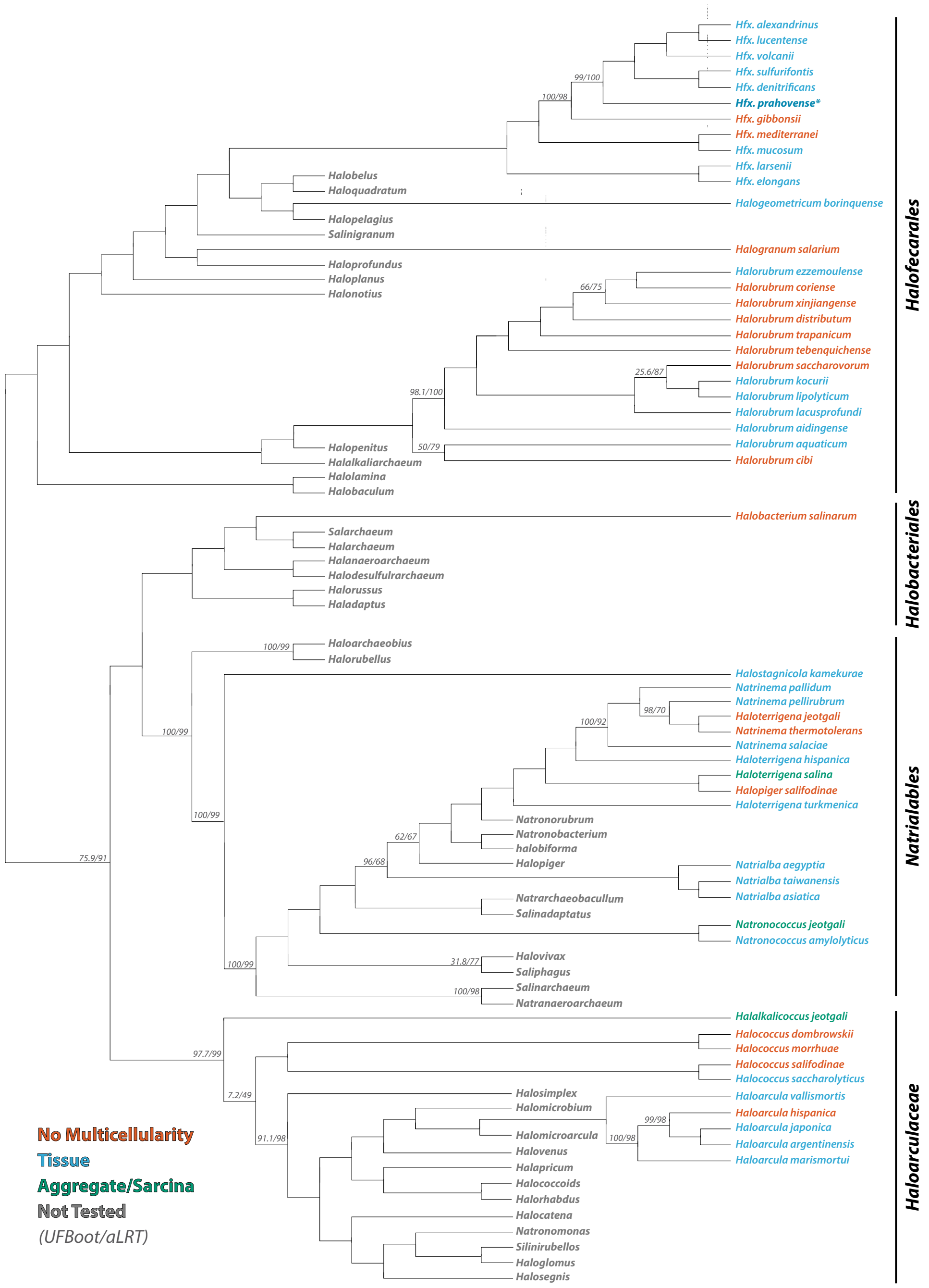


**Fig. S5. Evolutionary diversity of haloarchaeal tissues based on core genome.** The cladogram represents the evolutionary relationship between 315 Halobacteria genomes, estimated from the concatenation of 385 core genes (see Supplementary Methods for details on gene selection and alignment procedure). Ultrafast bootstrap approximation (UFBoot) and the approximate likelihood-ratio test (aLRT), used as bipartition support values, are shown for nodes that do not equal 100. Taxa are color-coded according to identified multicellular traits: red reflects taxa that multicellularity was not observed, blue denotes taxa with tissue organization, and green reflects taxa displaying sarcina-like aggregate structures. Collapsed taxa shown in grey have not been assessed in this study. For clarity in presentation, branch lengths are not depicted, hence the cladogram; and clades composed of not-evaluated taxa. These traits have been collapsed, indicated by a shorter tip length. Detailed information regarding tree topology, branch lengths, and full bipartition support values can be found in Supplementary Data S1.

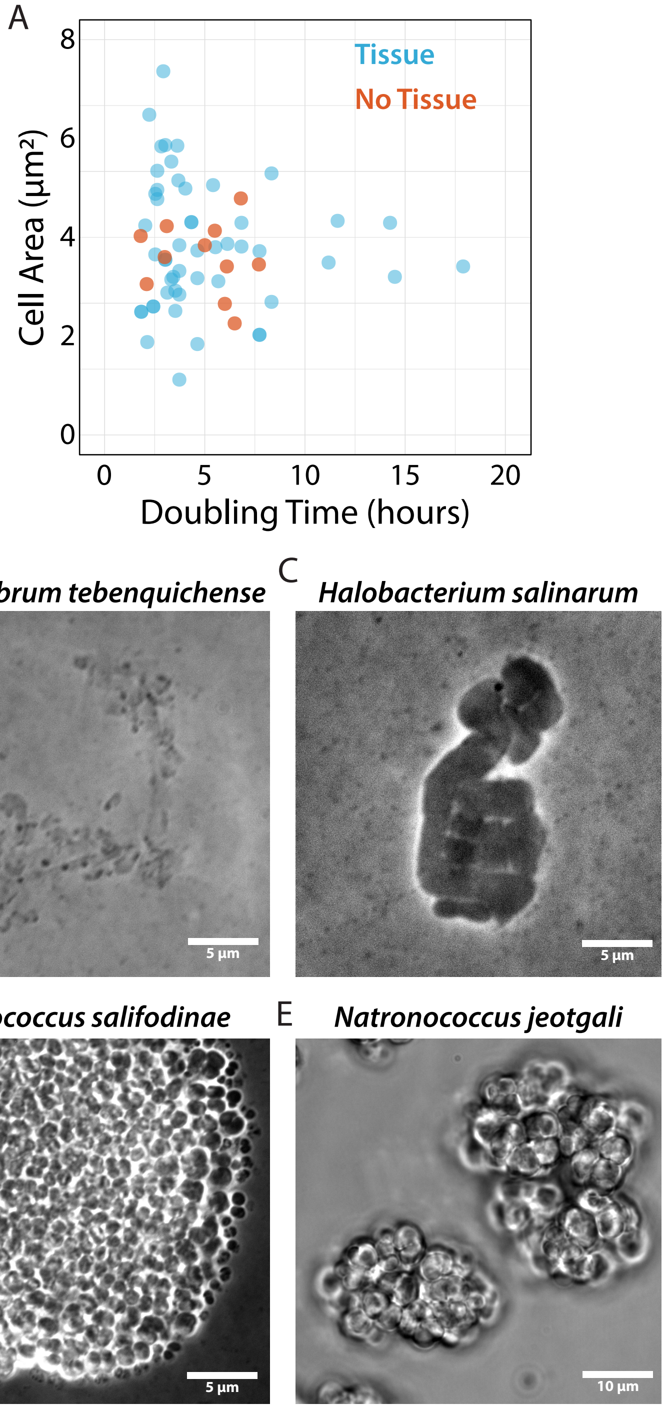


**Fig. S6: Diversity of responses other than tissue development of haloarchaeal cells to compression.** Phase contrast images of **(A)** *Halorubrum tebenquichense*, **(B)** *Halobacterium salinarum*, **(C)** *Halococcus salifodinae*, and **(D)** *Natronococcus jeotgali* under compression. **(E)** Correlation between unicell size (prior to compression) and doubling time from liquid cultures of tested haloarchaeal species.

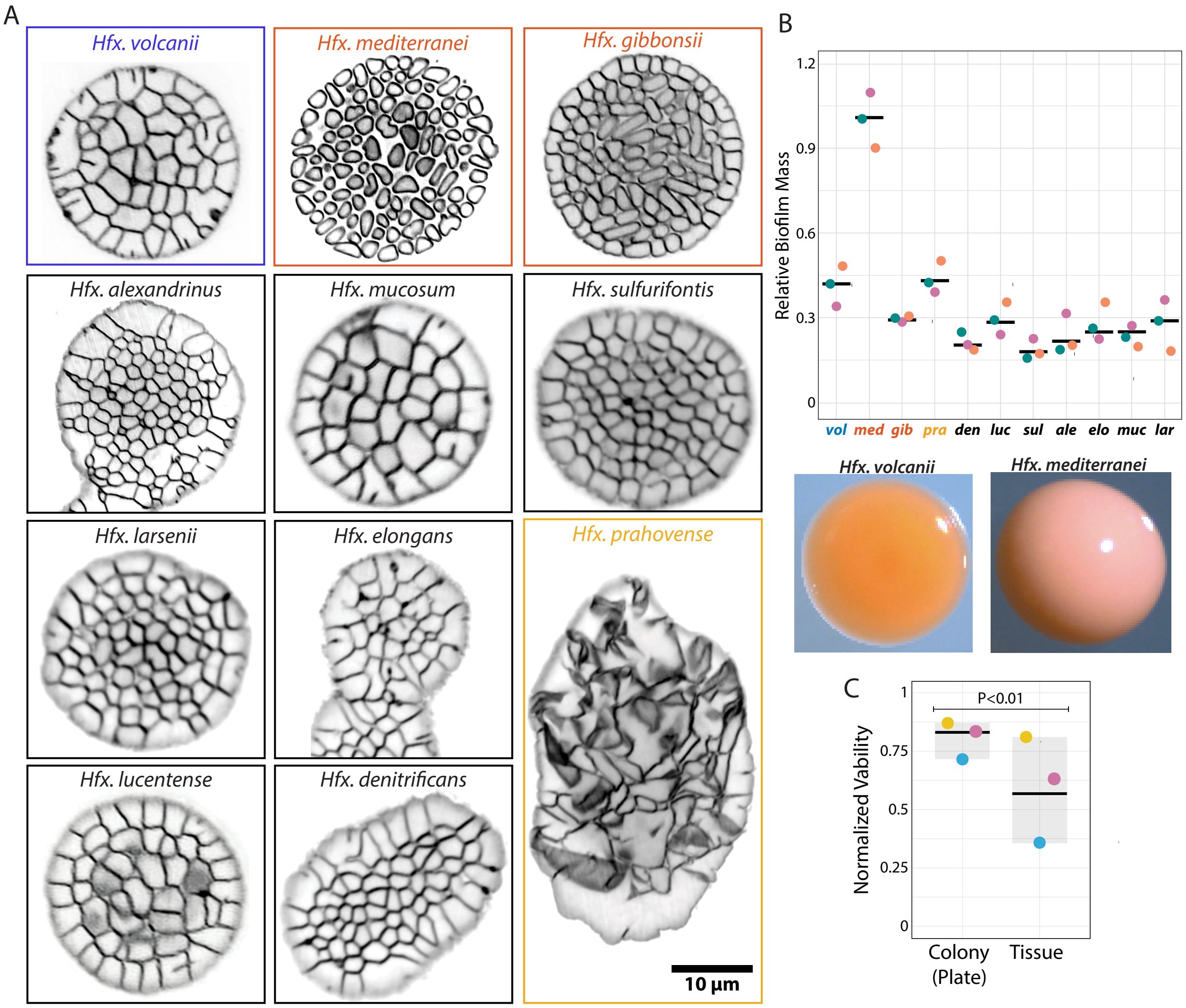


**Fig. S7. Diversity of archaeal tissue structures and biofilm production across *Haloferax* species. (A)** SoRa super-resolution microscopy images of 11 *Haloferax* species developed under compression. Tissues shown in *Hfx. volcanii* (blue frame). *Hfx. mediterranei* and *Hfx. gibbonsii* (red frames) cannot form tissues and *Hfx. prahovense* (yellow frame) form larger, deformed tissues. **(B)** (Top) Biofilm mass measurements from *Haloferax* species *Hfx. mediterranei* (med) displays markedly high biofilm formation relative to other tested *Haloferax* species. Representative photography of *Hvo* (bottom-left) and *Hmed* (bottom-right) colonies show the pale aspect of *Hmed* compared to *Hvo*. **(C)** Viability of *Hvo* tissues compared to *Hvo* cells from isolated colonies from plate streaks measured by colony formation unit (CFU). CFUs were normalized by liquid cultures used as initial inoculum. The center of the boxplots indicates the median and shades indicate the distribution range. Each datapoint represents a biological replicate.

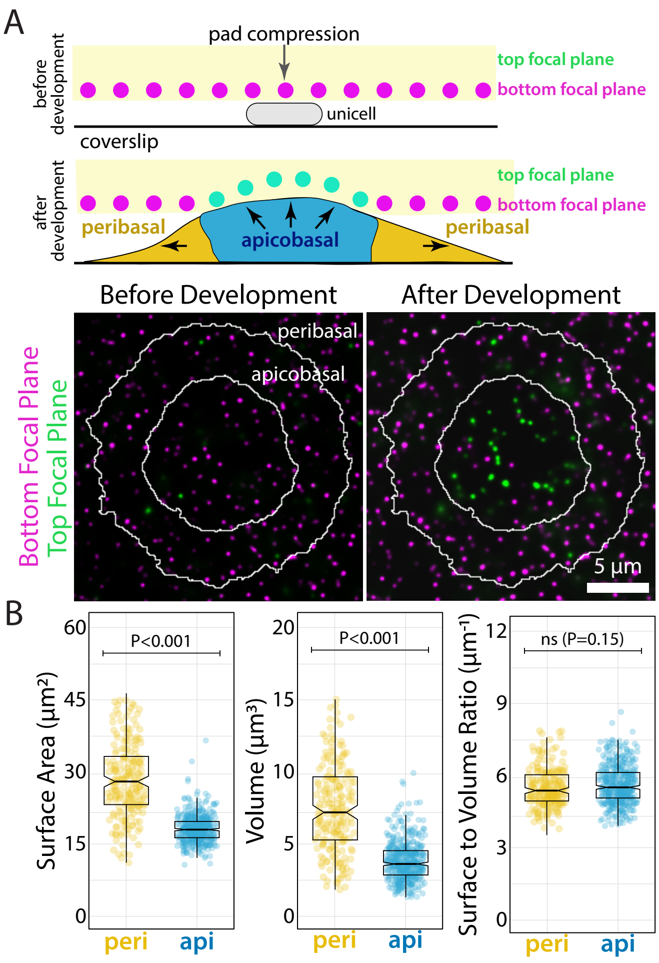


**Fig. S8. Apicobasal cells interact directly with pads, allowing peribasal cells to escape from mechanical compression. (A)** Traction force microscopy setup using fluorescent beads attached to the pad's surface compressing cells. Displacement of beads exclusively on top of apicobasal cells was captured by epifluorescence Z-stacks at the time of compression, and after 24 hours. Images were false colored in magenta (bottom focal plane) and green (top focal plane) **(B)** Surface area and volume measurements of apicobasal and peribasal cells from 3D projections.

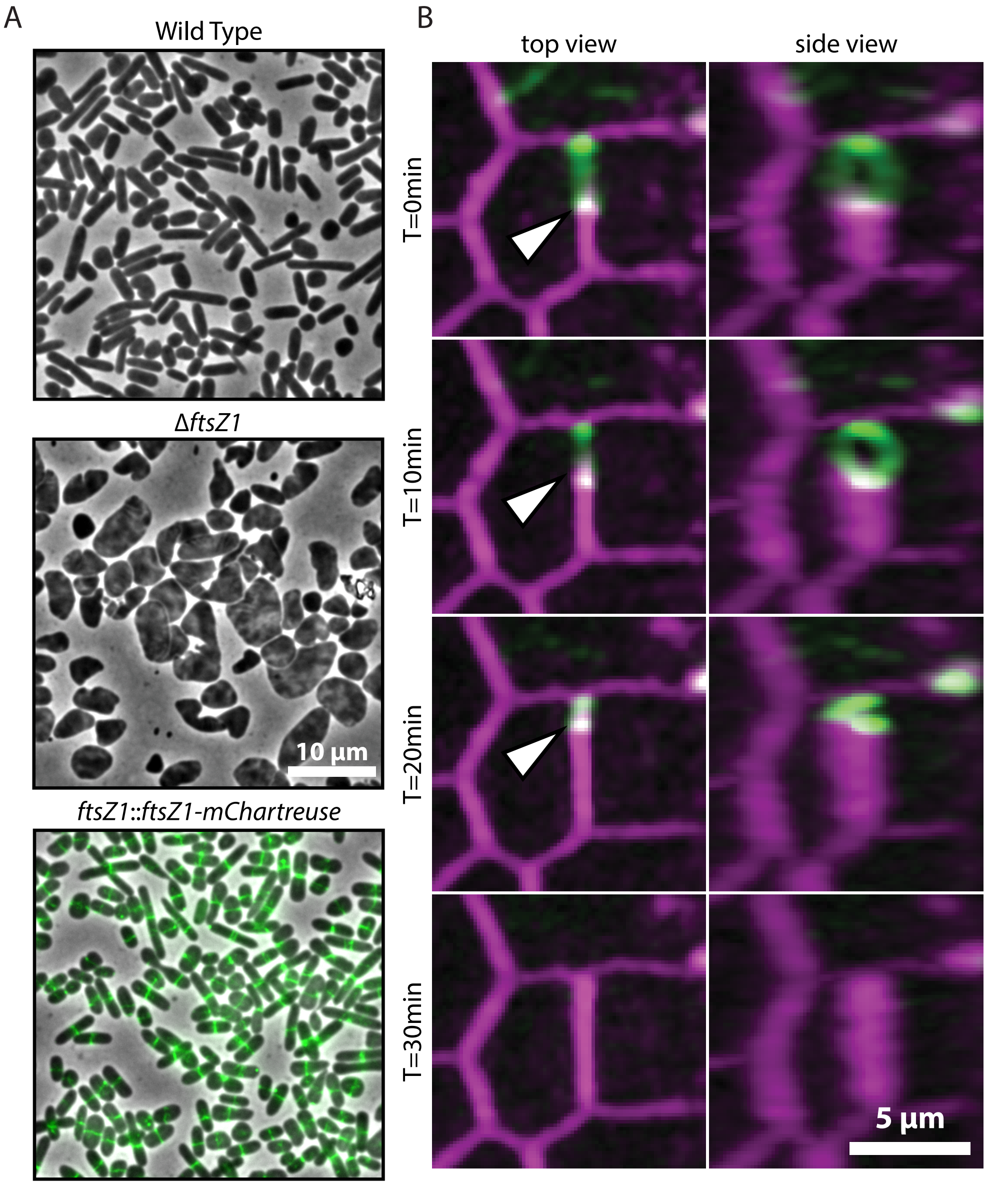


**Fig. S9. FtsZ1-mChartreuse is a functional fusion that localizes to the tissues' nascent junctions. (A)** Phase contrast images of (top) wild-type and (middle) Δ*ftsZ1* mutant cells. (bottom) Phase contrast and epifluorescence overlay of cells expressing FtsZ1-mChartreuse under the control of its native promoter. **(B)** SoRa super-resolution microscopy images of tissue junction formation from the top and side perspective of the 3D projection in 10-minute intervals (at 0, 10, 20, and 30 min). The white arrow indicates a nascent cell junction where the FtsZ1-mChartreuse (green) constricts together with the S-layer (magenta).

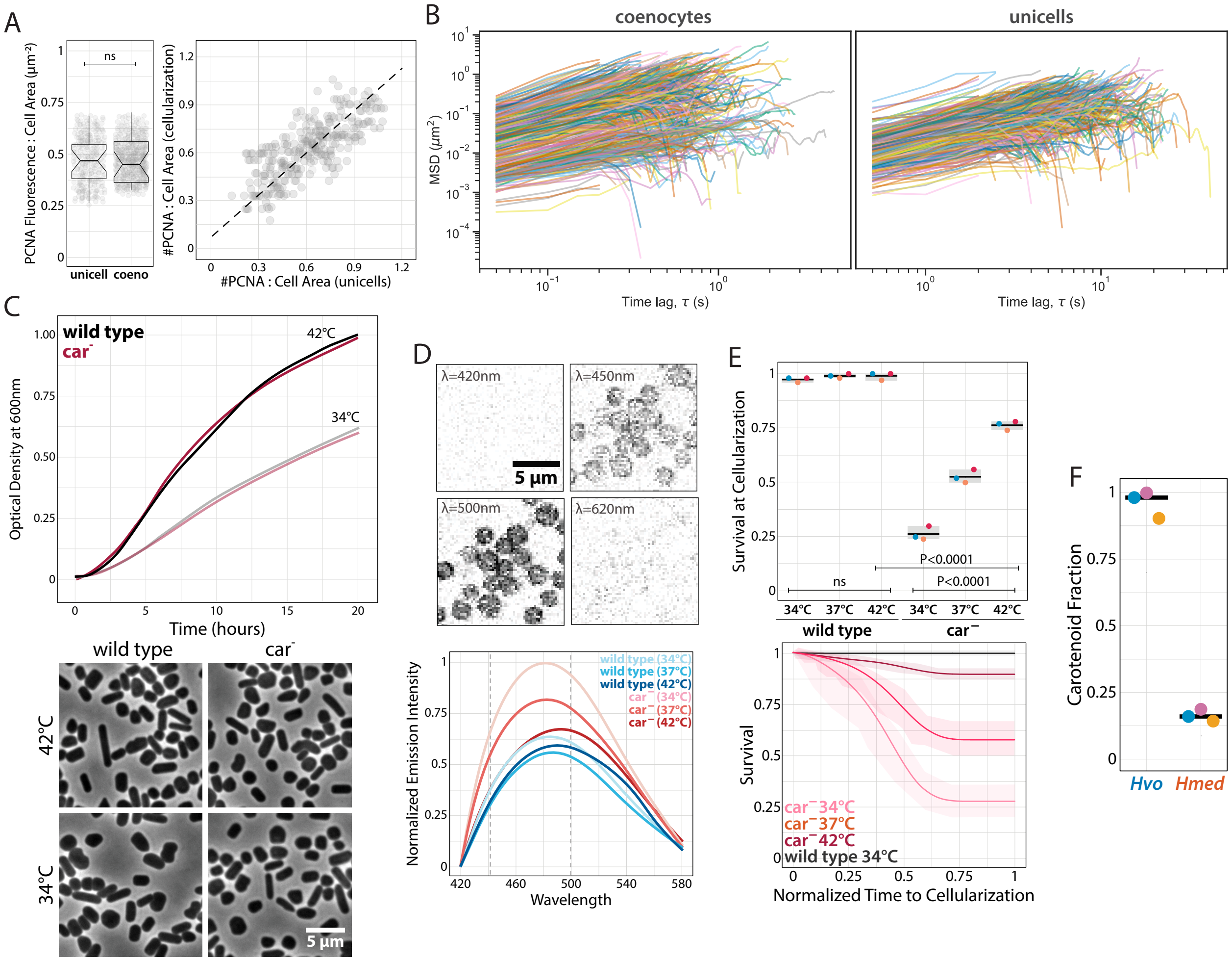

**Fig. S10. Membrane fluidity impacts viability during multicellular development. (A)** (Left) Sum of msfGFP-PCNA fluorescence intensity within foci in a cell normalized by cell area between unicells and coenocytes at the onset of cellularization. (Right) The number of msfGFP-PCNA foci in each cell normalized by cell area in unicells versus coenocytes at the onset of cellularization. **(B)** Mean Squared Displacement (MSD) vs time lag (τ) for individual full-length bSpoJ tracks ($R^2 \geq 0.9$) from unicells and coenocytes. Both axes are plotted with logarithmic scaling. **(C)** (Left) Growth curves of wild-type and car$^-$ liquid cultures at 42°C or 34°C. (Right) Phase contrast images of wild-type and car$^-$ unicells from liquid cultures at 42°C or 34°C. **(D)** (Left) Spectral confocal microscopy of representative wild-type spheroplasts stained with 100μM Laurdan at different wavelengths. (Right) Blue-shift emission spectra from confocal images of wild-type and car$^-$ cells at lower temperatures, indicating a decrease in membrane fluidity. **(E)** Normalized survival rates of wild-type and car- cells across temperatures. Survival was determined by cell lysis events from phase-contrast time-lapses. **(F)** Quantification of carotenoid production in *Hvo* and *Hmed*, showing higher production of pigments by *Hvo*.

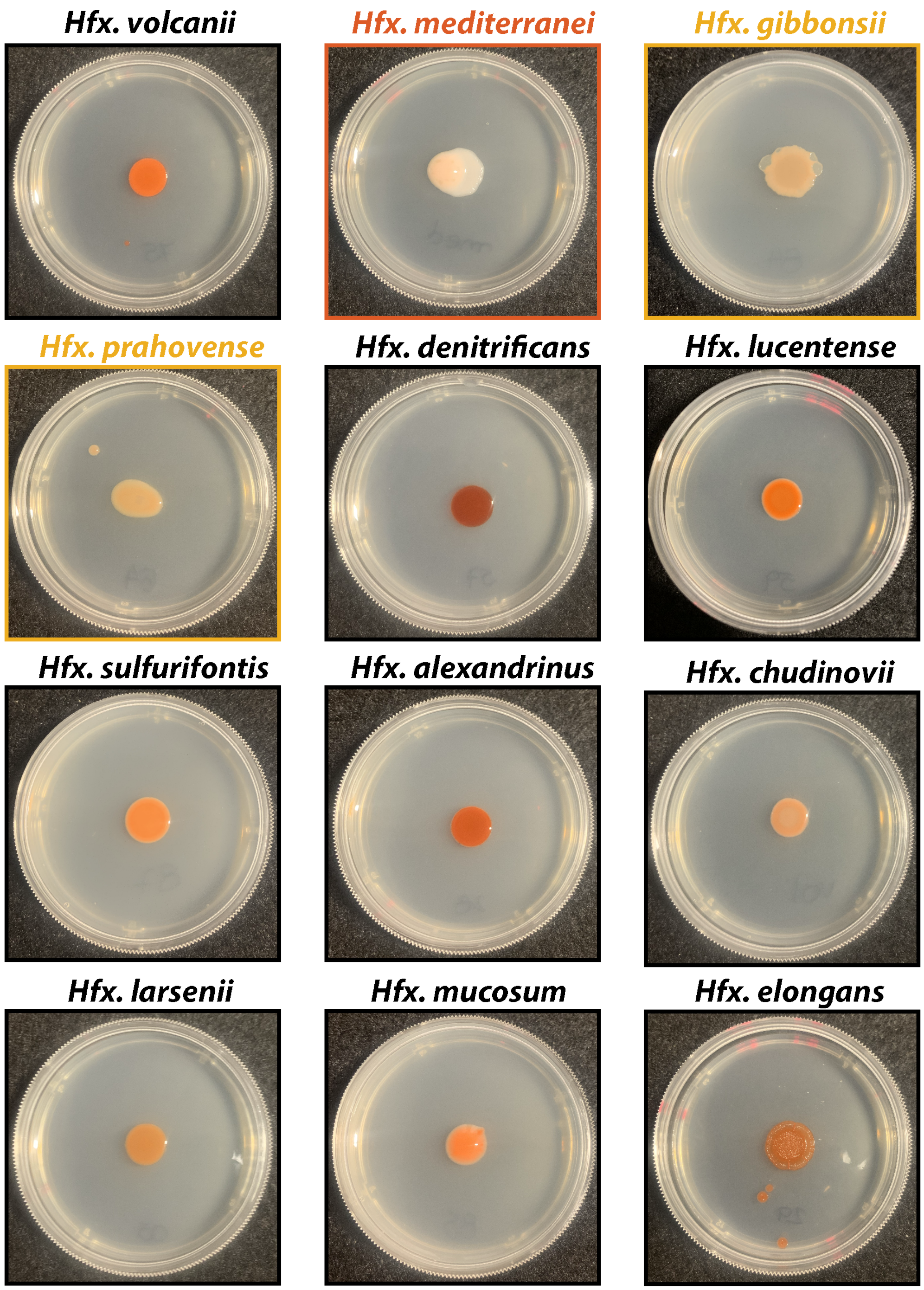


**Fig. S11. Archaeal tissue development correlates with carotenoid production across *Haloferax* species.** Colonies of twelve *Haloferax* strains on Hv-Cab agar plates. Colonies from species that do not develop into archaeal tissues (*Hfx. mediterranei*, red frame) have the lowest observed red coloration, whereas colonies from *Hfx. prahovense* and *Hfx. gibbonsii* (yellow frame), that form tissues with unstable junctions, show lower red coloration than other species.

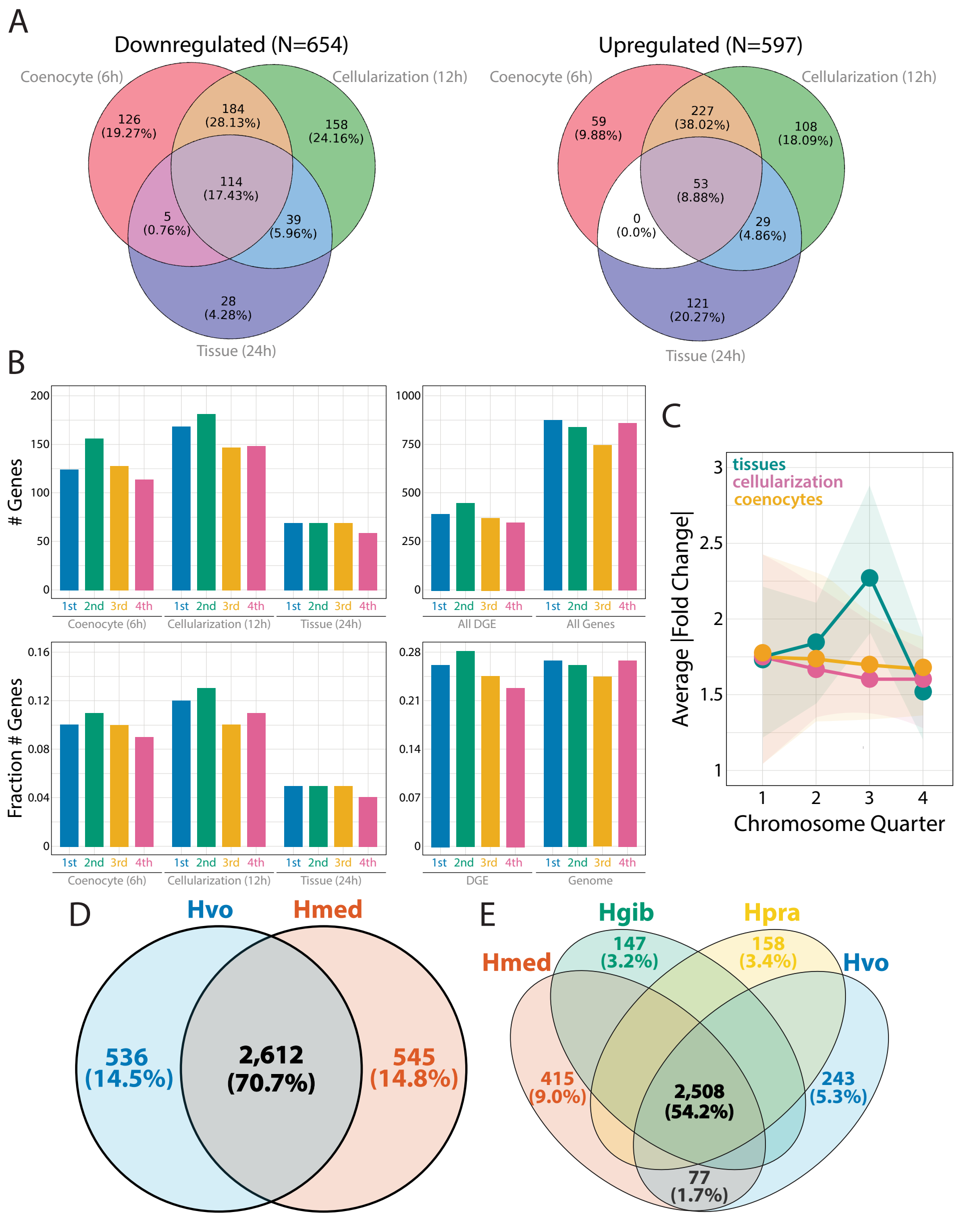


**Fig. S12. Temporal and genomic patterns observed from RNA-seq during Multicellular Development. (A)** Venn diagram with the number of downregulated (left) and upregulated (right) genes in each stage. **(B)** Number of differentially expressed gene candidates grouped by quadrant of the main chromosome of *Hvo* across each (top-left) or all (top-right) developmental stages. Number of differentially expressed gene candidates normalized by the number of total genes in each quadrant of the main chromosome of *Hvo* across each (bottom-left) or all (bottom-right) developmental stages. **(C)** The modulus of expression fold-change values from both upregulated and downregulated gene candidates mapped to each quadrant of the main chromosome of *Hvo*. A significant difference in the amplitude of gene expression was observed only in tissues, specifically in the third chromosomic quadrant. **(D)** Venn diagram of shared and exclusive ortholog distribution from comparative genomic analysis for *Hvo* (blue) and *Hmed* (orange). **(E)** Venn diagram of enrichment ortholog distribution between *Hvo* (blue), *Hmed* (orange), *Hfx. gibbonsii* (*Hgib*; green), and *Hfx. prahovense* (*Hpra*; yellow).

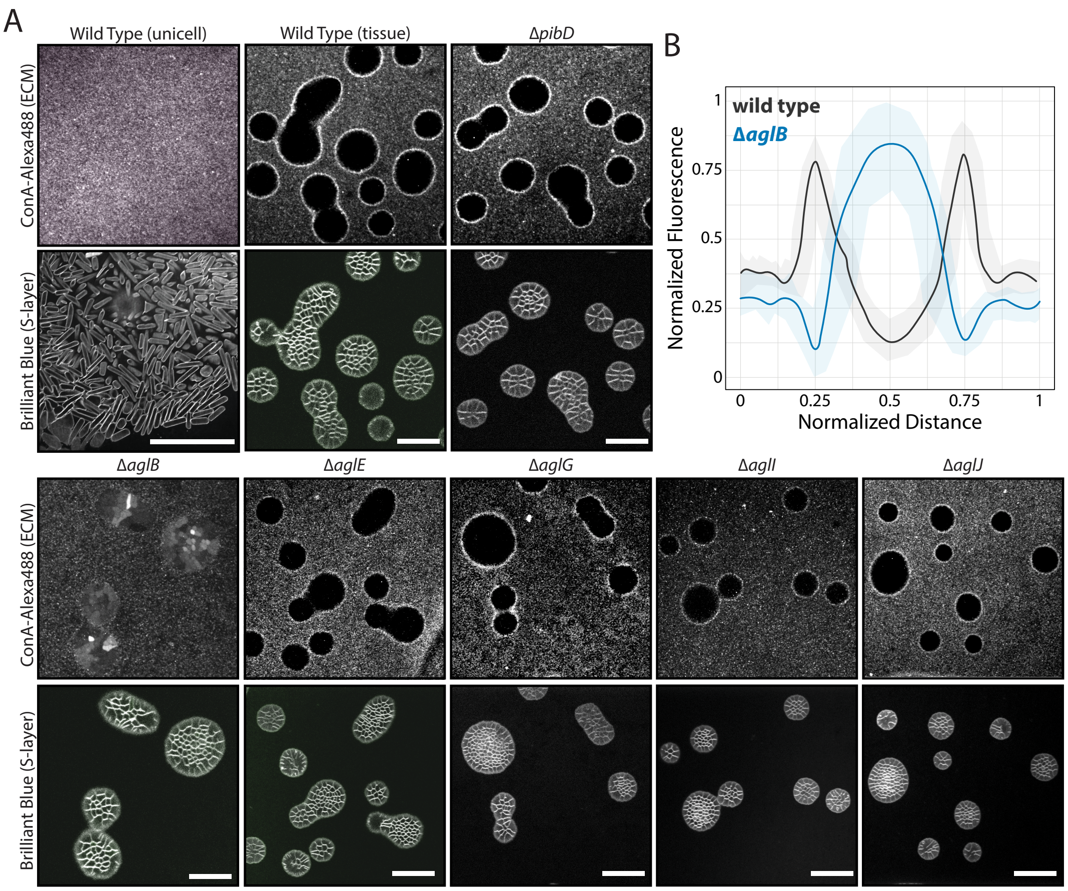


**Fig. S13. Polarity patterns of N-Glycosylation are tissue-specific and independent of biofilm. (A)** Spinning-disk confocal images of unicells and archaeal tissues across mutant strains stained with ConAAlexa488 **(B)** Normalized ConA-Alexa488 fluorescence intensity across wild-type (black) and Δ*aglB* (blue) tissues. Normalized distances across tissues were determined by the longest diameter measurement. Scale bars: 10 µm.

**Table S1.** ***Haloferax*** **species and RefSeq genome accessions used.**

| Species | RefSeq | Assembly | Forms Arc-Tissues |
|---|---|---|---|
| *Haloferax alexandrinus* | GCF_030535215.1 | ASM3053521v1 | Yes |
| *Haloferax denitrificans* | GCF_000337795.1 | ASM33779v1 | Yes |
| *Haloferax elongans* | GCF_000336755.1 | ASM33675v1 | Yes |
| *Haloferax gibbonsii* | GCF_000336775.1 | ASM33677v1 | **No** |
| *Haloferax larsenii* | GCF_000336955.1 | ASM33695v1 | Yes |
| *Haloferax lucentense* | GCF_000336795.1 | ASM33679v1 | Yes |
| *Haloferax mediterranei* | GCF_033843025.1 | ASM3384302v1 | **No** |
| *Haloferax mucosum* | GCF_000337815.1 | ASM33781v1 | Yes |
| *Haloferax prahovense* | GCF_000336815.1 | ASM33681v1 | ***Yes** |
| *Haloferax sulfurifontis* | GCF_014635105.1 | ASM1463510v1 | Yes |
| *Haloferax volcanii* | GCF_000025685.1 | ASM2568v1 | Yes |

*Cells cellularize into larger and unorganized tissues

**Table S2. Contingency table used in the hypergeometric test that allows for classification of proteins per class according to a given orthogroup.** N is the total protein population; K is the number of proteins issuing from the cohort of species that form tissue; n is the total number of proteins in an orthogroup; and k is the number of proteins from an orthogroup that issue from the group of species that form tissue.

| | **∈ orthogroup** | **∉ orthogroup** | **total** |
|---|---|---|---|
| **∈ sp. that forms tissue** | $k$ | $K - k$ | $K$ |
| **∉ sp. that forms tissue** | $n - k$ | $N + k - n - K$ | $N - K$ |
| **total** | $n$ | $N - n$ | $N$ |

**Table S3. Plasmids used in this work**

| Alias | Genotype | Reference |
|---|---|---|
| pTA962 | pTA962 (PtnA *pyrE2 hdrB ori-pHV2 oriC*) | (98) |
| pTA131 | pTA131 (*pyrE2*) | (98) |
| eTR68 | pTSD1 (*pyrE2*) | This work |
| eBL416 | pTDS1::ftsZ1-40aa-mChartreuse | This work |
| eSB14 | pTA131::mevR-HaloTag-15aa-mNG-40aa-PCNA | This work |
| eOL52 | pTA962::sec-spoIIIJ-HaloTag | This work |

**Table S4. Haloarchaeal strains used in this work**

| Alias | Genotype | Growth Media | Reference |
|---|---|---|---|
| ATCC 33960 | *Haloarcula hispanica* | 372-Cab | (99) |
| NRC-1 | *Halobacterium salinarum* | HCM | (100) |
| ATCC 33500 | *Haloferax mediterranei* | HV-Cab | (101) |
| DSM 4427 | *Haloferax gibbonsii* | HV-Cab | (99) |
| DSM 27206 | *Haloferax alexandrinus* | HV-Cab | (102) |
| DSM 4425 | *Haloferax denitrificans* | HV-Cab | (103) |
| DSM 14919 | *Haloferax lucentense* | HV-Cab | (104) |
| DSM 16227 | *Haloferax sulfurifontis* | HV-Cab | (105) |
| DSM 18310 | *Haloferax prahovense* | HV-Cab | (106) |
| DSM 26526 | *Haloferax chudinovii* | HV-Cab | (107) |
| DSM 27190 | *Haloferax larsenii* | HV-Cab | (108) |
| DSM 27191 | *Haloferax mucosum* | HV-Cab | (109) |
| DSM 27209 | *Haloferax elongans* | HV-Cab | (109) |
| DSM 11551 | *Halogeometricum borinquense* | 372-Cab | (110) |
| DSM 23171 | *Halogranum salarium* | HV-Cab | (111) |
| DSM 17463 | *Halorubrum ezzemoulense* | 372-Cab | (112) |
| DSM 10284 | *Halorubrum coriense* | 372-Cab | (113) |
| DSM 21996 | *Halorubrum xinjiangense* | 372-Cab | (114) |
| DSM 23453 | *Halorubrum distributum* | 372-Cab | (115) |
| DSM 12287 | *Halorubrum trapanicum* | 372-Cab | (116) |

| | | | |
|---|---|---|---|
| DSM 14210 | *Halorubrum tebenquichense* | 372-Cab | (117) |
| DSM 1137 | *Halorubrum saccharovorum* | 372-Cab | (118) |
| DSM 19804 | *Halorubrum kocurii* | 372-Cab | (119) |
| DSM 21995 | *Halorubrum lipolyticum* | 372-Cab | (120) |
| DSM 5036 | *Halorubrum lacusprofundi* | 372-Cab | (121) |
| DSM 23496 | *Halorubrum aidingense* | 372-Cab | (120) |
| DSM 18322 | *Halorubrum aquaticum* | 372-Cab | (122) |
| DSM 19504 | *Halorubrum cibi* | HV-Cab | (123) |
| DSM 22472 | *Halostagnicola kamekurae* | 372-Cab | (124) |
| DSM 3751 | *Natrinema pallidum* | 372-Cab | (125) |
| DSM 15624 | *Natrinema pellirubrum* | 372-Cab | (125) |
| DSM 18796 | *Halalkalicoccus jeotgali* | 372-Cab | (126) |
| DSM 18795 | *Natronococcus jeotgali* | 372-Cab | (127) |
| DSM 18794 | *Haloterrigena jeotgali* | 372-Cab | (128) |
| DSM 11552 | *Haloterrigena thermotolerans* | 372-Cab | (129) |
| DSM 25055 | *Natrinema salaciae* | 372-Cab | (130) |
| DSM 18328 | *Haloterrigena hispanica* | 372-Cab | (131) |
| DSM 26231 | *Halopiger salifodinae* | 372-Cab | (132) |
| DSM 5511 | *Haloterrigena turkmenica* | 372-Cab | (133) |
| DSM 13077 | *Natrialba aegyptia* | 372-Cab | (134) |
| DSM 12281 | *Natrialba taiwanensis* | 372-Cab | (134) |

| | | | |
|---|---|---|---|
| DSM 12278 | *Natrialba asiatica* | 372-Cab | (135) |
| DSM 10524 | *Natronococcus amylolyticus* | 372-Cab | (136) |
| DSM 14522 | *Halococcus dombrowskii* | 372-Cab | (137) |
| DSM 1307 | *Halococcus morrhuae* | 372-Cab | (138) |
| DSM 8989 | *Halococcus salifodinae* | 372-Cab | (139) |
| DSM 5350 | *Halococcus saccharolyticus* | 372-Cab | (140) |
| DSM 3756 | *Haloarcula vallismortis* | 372-Cab | (141) |
| DSM 6131 | *Haloarcula japonica* | 372-Cab | (142) |
| DSM 12282 | *Haloarcula argentinensis* | 372-Cab | (143) |
| DSM 3752 | *Haloarcula marismortui* | 372-Cab | (144) |

**Table S5.** ***Haloferax volcanii*** **strains used in this work**

| Alias | Genotype | Reference |
|---|---|---|
| DS2 | *Haloferax volcanii* | (145) |
| H26 | Δ*pyrE2* | (146) |
| H53 | Δ*pyrE2* Δ*trpA* | (146) |
| aBL126 | Δ*pyrE2 volA::volA-40aa-msfGFP-pyrE2* | (43) |
| aKA17 | Δ*pyrE2* Δ*volA* | (44) |
| ID112 | Δ*hdrB* Δ*pyrE2* Δ*ftsZ1* Δ*ftsZ2* | (147) |
| ID77 | Δ*pyrE2* Δ*ftsZ2* | (147) |
| Δ2528 | Δ*pyrE2* Δ*HVO_2528* | (148) |
| MT4 | Δ*pibD* | (149) |
| ΔaglB | Δ*pyrE2* Δ*trpA tn::aglB* | (150) |
| ΔaglE | Δ*pyrE2* Δ*trpA tn::aglE* | (150) |
| ΔaglJ | Δ*pyrE2* Δ*trpA tn::aglJ* | (151) |
| ΔaglG | Δ*pyrE2* Δ*trpA tn::aglG* | (151) |
| ΔaglI | Δ*pyrE2* Δ*trpA tn::aglI* | (152) |
| aBL582 | Δ*pyrE2* ftsZ1::ftsZ1-40aa-mChartreuse | This work |
| aSB14 | *pcna::mevR-HaloTag-15aa-mNG-40aa-pcna* | This work |
| aOL39 | Δ*pyrE2* pTA962::sec-spoIIIJ(TM)-HaloTag | This work |

**Table S6. Oligos used in this work**

| Alias | Sequence (5' → 3') |
|---|---|
| oHV13 | CGGACCTATTGCGCATATGACAAAGCTCAAAGATCAAACG |
| oAB130 | GCCTTGACCTGGGCCAGATC |
| oBL97 | AGGTGGCACTTTTCGG |
| oBL105 | TGAGCAAAAGGCCAGC |
| oBL318 | GGAATTCGATATCAAGCTTATCGATTTTTAGCCGCTGATTTCTAAGGTAG |
| oBL354 | CTTGAGGGTAGCGGAC |
| oBL532 | GATATCGAATTCCTGCAGCCTCGAGCCGTCCCG |
| oBL533 | CGAGGGGTTTTATCCACGCGAAGAAACGGTTTTGTGG |
| oBL536 | GAAAAGTGCCACCTGAATTCACATGCATGGGGGGTTG |
| oBL537 | GGGTTTTATCCACGGAATTCCATGAGCTTCTTTGATTCGAGC |
| oOL79 | GGCTCTTTCACGAACGATTCGCTGTCAG |
| oOL80 | GTGAAAGAGCCGATCACTG |
| oOL81 | GCTACCCTCAAGCTTTTTCTTTCCTCCGGCTTTT |
| oTR202 | CCGAAAAGTGCCACCTCGGCCACGCGGTC |
| oTR203 | CCTACGACGCGTGAGGTATGGCGCGCCGGTCCGAG |
| oTR204 | CGCCATACCTCACGC |
| oTR205 | CGAGGGGTTTTATCCACGGCCCGGTTCGGGT |
| oSB41 | CCGAAAAGTGCCACCTCGTGACGCTCGCG |
| oSB42 | CCATCCCCTCCCATGATGTGCCGTCGTACG |

**Table S7. Statistical Information from Datasets in this Study.**

| Figure | Experiment | N Samples (Biological Replicate) | Statistical Test |
|---|---|---|---|
| 1G | Unicell Ablation Displacement | 119 (14,48,67) | Kolmogorov-Smirnov |
| | Tissue Ablation Displacement | 92 (12,42,38) | |
| 1H | Unicell Ablation Junction Recoil | 22 (6,13,3) | - |
| | Tissue Ablation Junction Recoil | 24 (4,12,6) | |
| 2E | *Hvo* x Hmed Survival | 3 (1,1,1) | One-Way ANOVA |
| 3D | Unicell Growth Rate | 942 (333,278,341) | Kolmogorov-Smirnov |
| | Peribasal Growth Rate | 172 (56,74,42) | |
| | Apicobasal Growth Rate | 205 (99,63,43) | |
| 3E | Unicell Lifespan | 896 (211,405,280) | Kolmogorov-Smirnov |
| | Peribasal Lifespan | 99 (52,25,22) | |
| | Apicobasal Lifespan | 191 (87,40,64) | |
| 3F | Unicell Deformation | 337 (109,58,170) | Kolmogorov-Smirnov |
| | Peribasal Deformation | 46 (23,9,14) | |
| | Apicobasal Lifespan | 121 (25,82,14) | |
| 4C | WT Cellularization | 241 (73,90,78) | Kolmogorov-Smirnov |
| | ΔftsZ2 Cellularization | 164 (33,69,62) | |
| 4D | Unicell bSpoJ Diffusion | 1782 (1023,455,304) | Kolmogorov-Smirnov |
| | Coenocyte bSpoJ bSpoJ Diffusion | 1875 (521,179,1175) | |
| 4E | WT Generalized Polarization | 1455 (654,324,477) | - |
| | Car$^-$ Generalized Polarization | 1199 (560,340,299) | |
| 4F | WT Cellularization Area at 34°C | 106 (69,26,11) | Kolmogorov-Smirnov |
| | WT Cellularization Area at 37°C | 131 (45,24,52) | |
| | WT Cellularization Area at 42°C | 162 (39,102,21) | |
| | Car$^-$ Cellularization Area at 34°C | 77 (26,19,32) | |
| | Car$^-$ Cellularization Area at 37°C | 125 (59,23,43) | |

| | | | |
|---|---|---|---|
| | Car$^-$ Cellularization Area at 42°C | 179 (91,26,62) | |
| 4G | WT Unicell bSpoJ Diffusion | 2522 (1306,560,656) | Kolmogorov-Smirnov |
| | WT Coenocyte bSpoJ Diffusion | 1965 (1048,753,164) | |
| | WT Cellularization bSpoJ Diffusion | 1358 (509,515,334) | |
| | Car$^-$ Unicell bSpoJ Diffusion | 2114 (734,251,1129) | |
| | Car$^-$ Coenocyte bSpoJ Diffusion | 987 (735,201,51) | |
| | Car$^-$ Cellularization bSpoJ Diffusion | 793 (282,405,106) | |
| 5B | Volactin-GFP Unicell Fluorescence | 103 (21,44,38) | - |
| | Volactin-GFP Tissue Fluorescence | 42 (9,16,17) | |
| | GFP Tissue Fluorescence | 35 (10,5,20) | |
| 5D | Volactin Unicell Cable Angle | 172 (41,62,71) | Kolmogorov-Smirnov |
| | Volactin Coenocyte Cable Angle | 94 (33,16,45) | |
| | Volactin Tissue Cable Angle | 79 (18,32,29) | |
| 5E | WT Coenocyte Height:Area Ratio | 152 (43,67,42) | Kolmogorov-Smirnov |
| | Δ*volA* Coenocyte Height:Area Ratio | 117 (22,52,43) | |
| S1B | Cell Solidity (Liquid Bulk Culture) | 931 (255,245,431) | - |
| | Cell Solidity (Microfluidics) | 1525 (350, 500, 675) | |
| | Cell Solidity (0.25% Agarose Pad) | 134 (11,56,67) | |
| | Cell Solidity (0.5% Agarose Pad) | 102 (34,23,45) | |
| | Cell Solidity (1% Agarose Pad) | 91 (20,42,39) | |
| S1C | DMA Agarose Pads | 3 (1,1,1) | - |
| S2C | Cellularization Cell Area (2% Agarose Pad) | 211 (50,50,111) | Kolmogorov-Smirnov |
| | Cellularization Cell Area (2.5% Agarose Pad) | 198 (50,50,98) | |
| | Cellularization Cell Area (3% Agarose Pad) | 185 (50,50,85) | |
| | Cellularization Cell Area (3.5% Agarose Pad) | 201 (50,50,101) | |

| | | | |
|---|---|---|---|
| S7B | *Hfx* Species Biofilm Mass | 3 (1,1,1) | One-Way ANOVA |
| S7C | *Hvo* Colonies x Tissues Survival | 3 (1,1,1) | One-Way ANOVA |
| S8B | Peribasal Cell Surface:Area | 120 (63,32,25) | Kolmogorov-Smirnov |
| | Apicobasal Cell Surface:Area | 230 (77,64,89) | |
| S10A | Unicell PCNA Fluorescence | 158 (42,51,55) | Kolmogorov-Smirnov |
| | Cellularization PCNA Fluorescence | 125 (37,65,23) | |
| S10C | WT x Car$^-$ Growth Curves | 3 (1,1,1) | - |
| S10E | WT Coenocyte Survival at 34°C | 154 (67,42,35) | Kolmogorov-Smirnov |
| | Car$^-$ Coenocyte Survival at 34°C | 91 (30,50,11) | |
| | Car$^-$ Coenocyte Survival at 37°C | 138 (25,39,74) | |
| | Car$^-$ Coenocyte Survival at 42°C | 112 (54,33,35) | |
| S10F | *Hvo* x *Hmed* Carotenoid Fraction | 3 (1,1,1) | - |
| S13B | WT ConA-Alex488 Localization | 42 (12,21,9) | - |
| | $\Delta$*aglB* ConA-Alex488 Localization | 61 (31,19,11) | |

**Movie S1. *Hfx. volcanii* unicell growth within ArcCell microfluidics and under agarose pads**. Phase-contrast microscopy timelapses in ArcCell (left panel) and under 0.25% agarose pads (right panel). Timelapses were acquired at 2- and 5-min intervals, respectively. Movies are displayed at 8 frames per second. Scale bars: 5µm

**Movie S2. Archaeal tissue development under compression.** Epifluorescence (left panel) and phase-contrast (right panel) microscopy of cells growing under compression by a 2.5% Hv-Cab agarose pads. Timelapses collected at 20-min intervals and displayed at 5 frames per second. S-layer stained with Brilliant Blue. Scale bar: 5µm

**Movie S3. DNA replication carries on during the coenocytic phase.** Epifluorescence microscopy of mNeonGreen-PCNA (DNA sliding clamp, cyan) foci decorating replication sites. Middle panel: phase contrast. Right panel: mNeonGreen-PCNA. Left panel: overlay. Timelapses were collected at 2-min intervals and displayed at 20 frames per second. Scale bar: 5µm

**Movie S4. 3D-SoRa super-resolution microscopy of tissues versus unicellular colonies.** 3D projections from the central regions of a clustered unicell colony (left panel) and tissue (right panel). Z-stacks were collected at 0.1-µm slices. S-layer stained with Brilliant Blue. Scale bar: 5µm.

**Movie S5. Wounding of archaeal tissues induces directional stretching of cell junctions towards the injured site.** Central regions of unicell clusters (left panel) and tissues (right panel) were ablated by a 405 nm laser pulse (darker). Following injury, samples were imaged by SoRa super-resolution microscopy at 2.5-min intervals. For the tissue movie, a Z-stack projection with 2 slices separated by 0.5 µm is shown. Movies are displayed at 5 frames per second. S-layer stained with Brilliant Blue. Scale bar: 5µm.

**Movie S6. Cell membrane recoil of unicells and tissues by laser ablation.** Cytoplasmic membrane of unicells (left panel) and junctions within archaeal tissues (right panel) ablated with 405nm laser pulse. Following severing, samples were imaged by SoRa super-resolution microscopy at 1-second (unicells) and 100-ms (tissues) intervals. Movies are displayed at 10 frames per second. S-layer stained with Brilliant Blue. Scale bar: 5µm.

**Movie S7. *Hfx. mediterranei* cells do not form tissues and "swarms out" out compression zones.** Phase-contrast microscopy of *Hmed* cells growing under 3% (left panel) and 5% (right panel) agarose pads were collected at 15-min intervals. Movies are displayed at 5 frames per second. Scale bar: 5µm.

**Movie S8. *Hfx.* cells growth under microfabricated "escape-room" pillar traps.** Phase-contrast imaging of *Hvo* (right panel) and *Hmed* (left panel) cells were acquired at 10- and 1-min intervals, respectively. Movies are displayed at 5 frames per second. Scale bars: 10µm.

**Movie S9. Reversible Rod-shape transition after recovery from tissue shear-fragmentation.** (Left Panel) Phase-contrast timelapse of tissues being shear-shocked by a liquid wave (false-colored in magenta), followed by fast development (under a generation time) to swimming rods. (Right Panel) Zoom-in of a second tissue structure transition to rods followed by fragmentation. Movies were collected at 1-min intervals and displayed at 10 frames per second. Scale bar: 10µm.

**Movie S10. Super-resolution microscopy of peribasal and apicobasal cells.** (Top-Left Panel) 3D-STED projection of cells stained with Brilliant Blue acquired with 75-nm spaced Z slices. (Top-Right

Panel) 3D outline masks of apicobasal scutoid cells segmented from 3D-STED projections. (Bottom panels) 3D-iSIM projection of cells constitutively expressing cytoplasmic msfGFP acquired with 200-nm spaced Z slices. Scale bars: 5µm.

**Movie S11. Tissue recompression followed by shear-fragmentation.** Phase-contrast timelapse of tissues being disrupted by liquid shear-shock, followed by removal of liquid media and recompression of fragmented cells. Movies were collected at 1-min intervals and displayed at 20 frames per second. Scale bar: 10µm.

**Movie S12. Single-molecule tracking of the membrane-associated chimera bSpoJ-JFx549 conjugates in coenocytes.** Total Internal Reflection Fluorescence Microscopy (TIRFM) of coenocytes (shown in phase contrast) expressing bSpoJ conjugated with JFx549 dyes imaged at 250-millisecond intervals. Movies are displayed at 40 frames per second. Tracks (right panel) are false colored in a mean speed scale between 0.1
(blue) and 2 (red) µm/s. Scale bar: 5µm.

**Movie S13. Single-molecule tracking of the membrane-associated chimera bSpoJ-JFx549 conjugates in unicells.** Total Internal Reflection Fluorescence Microscopy (TIRFM) of unicells (shown in phase contrast) expressing bSpoJ conjugated with JFx549 dyes imaged at 500-ms intervals. Movies are displayed at 10 frames per second. Tracks (right panel) are false colored in a mean speed scale between 0.1 (blue) and 2 (red) µm/s. Scale bar: 5µm.

**Movie S14. Volactin endogenous expression increases as cells differentiate into tissues.**
Epifluorescence microscopy of volA-msfGFP under native promoter shows increase in fluorescence signal as cells differentiate into tissues. S-layer stained with Brilliant Blue. Scale bar: 5µm.

**Movie S15. Volactin dynamics in tissues.** Epifluorescence microscopy of volA-msfGFP acquired at 25-min intervals and displayed at 4 frames per second. Scale bar: 5µm.

**Movie S16. Volactin cables relative orientation during development.** 3D-SoRa super-resolution representative projections of volA-msfGFP cables in unicells (first/left panel), coenocytes (second panel), coenocytes during cellularization (third panel), and tissues (fourth/right panel). S-layer stained with Brilliant Blue. Z-stacks collected under 0.1-µm intervals. Scale bars: 5µm.

**Movie S17. Protein Glycosylation patterns in wild-type and Δ*aglB* tissues**. 3D projections of tissues showing S-layer (left panel, black) and protein glycosylation (right panel, blue) spatial profiles. Spinning-disk confocal Z-stacks were collected under 0.1-µm intervals. Glycoproteins are stained with ConA-Alexa 488. S-layer is stained with Brilliant Blue. Scale bars: 5µm.

**Data S1. Maximum likelihood phylogenetic tree of Haloarchaea with branch support values (JSONformatted TreeFile/FigTree).**

**Data S2. Differentially gene expression (DGE) candidates across developmental stages (XLSXformatted spreadsheet).** (A) All candidates above arbitrary threshold. (B) Shortlisted genes with possible role in development. Blue: upregulated. Orange: downregulated.

**Data S3. *Hfx. volcanii* unique genes compared to *Hfx. mediterranei* (XLSX-formatted spreadsheet).**

**Data S4. *Hfx. volcanii* unique genes compared to *Hfx. mediterranei, Hfx. gibbonsii, and Hfx. prahovense* (XLSX-formatted spreadsheet).**

**Data S5. Enrichment of orthologs among Hfx species that form tissues relative to *Hfx. mediterranei, Hfx. gibbonsii, and Hfx. prahovense* (XLSX-formatted spreadsheet)**
